# Profiling Germline Adaptive Immune Receptor Repertoire with gAIRR Suite

**DOI:** 10.1101/2020.11.27.399857

**Authors:** Mao-Jan Lin, Yu-Chun Lin, Nae-Chyun Chen, Allen Chilun Luo, Sheng-Kai Lai, Chia-Lang Hsu, Jacob Shujui Hsu, Chien-Yu Chen, Wei-Shiung Yang, Pei-Lung Chen

**Author notes:** these authors contributed equally to this work (Mao-Jan Lin, Yu-Chun Lin, Nae-Chyun Chen).

## Abstract

Genetic profiling of germline adaptive immune receptor repertoire (AIRR), including T cell receptor (TR) and immunoglobulin (IG), is imaginably relevant to numerous immune-related conditions, but currently insurmountable due to high genetic complexity. Our gAIRR Suite comprises three modules. gAIRR-seq, a probe capture-based targeted sequencing pipeline, profiles AIRR from individual DNA samples. gAIRR-call and gAIRR-annotate call alleles from gAIRR-seq reads and annotate whole-genome assemblies respectively. We gAIRR-seqed TRV and TRJ of seven Genome in a Bottle (GIAB) DNA samples with 100% accuracy, and discovered novel alleles. We also gAIRR-seqed and gAIRR-called a subject from both the peripheral blood mononuclear cells (PBMC) and oral mucosal cells. The calling results from these two cell types have a high concordance (99% for all known AIRR alleles). We gAIRR-annotated 36 genomes to cumulatively unearth 325 novel TRV alleles and 29 novel TRJ alleles. We could further profile the flanking sequences, including the recombination signal sequence (RSS). We validated two structural variants for HG002. We uncovered substantial conflicts of AIRR genes in references GRCh37 and GRCh38. The gAIRR Suite can potentially benefit future genetic study and clinical applications of various immune-related phenotypes.

## Introduction

Germline AIRR is the collection of TR and IG, which are composed of variable (V), diversity (D) and joining (J) regions. The receptors’ final sequences require somatic V(D)J recombination, where one gene from each V, (D) and J genetic region are selected and rearranged into a continuous fragment^1,2^, with or without additional somatic hypermutation. Although humans’ immune repertoire has theoretically extreme huge variability capable of binding all kinds of antigens^3–5^, for each individual the number of V(D)J genes/alleles in one’s genome is limited^3^. Therefore, any given individual’s immunological response may be restricted according to the germline AIRR composition^3,6,7^. Early data suggested SARS-CoV-2 antibodies prefer to use certain IG genes/alleles^8^. Carbamazepine-induced Stevens-Johnson syndrome^9^, besides the well-known *HLA-B*15:02* susceptibility in Asians, also depends on specific public TRs for the severe cutaneous adverse reaction^10^. In a simian immunodeficiency virus infection model, the TR alleles responsible for the potent CD8+ T cell response were recently determined^11^. Germline AIRR alleles have started to be found associated with human diseases, including rheumatic heart disease^12^, and Kawasaki disease^13,14^. These examples are likely only the tip of the iceberg because of the complexity in determining individual germline AIRR and the lack of diverse and comprehensive AIRR database^15^.The international ImMunoGeneTics (IMGT) database^5^, the central depository of germline AIRR, is a valuable resource but has been slow in adding new alleles.

Both genomic DNA (gDNA) and messenger RNA (mRNA)/complementary DNA (cDNA) can be used to profile AIRR of an individual, albeit yielding quite different information^16,17^. The popular mRNA/cDNA approach provides useful dynamic information of AIRR clonotypes and expression levels, which can be valuable if, and only if, the most relevant lymphocytes can be retrieved from the right site at the right time^16^. mRNA/cDNA-based AIRR sequencing at any given time point can only cover part of the full spectrum of germline AIRR. Depending on the library preparation strategies, the mRNA/cDNA approach might suffer from allele dropout, lack of full-length coverage and inconsistent results^18^. Furthermore, the non-coding flanking sequences, including RSS which may significantly affect recombination efficiency, cannot be profiled in the mRNA/cDNA approach. On the other hand, bulk gDNA includes non-rearranged and rearranged V(D)J segments and preserves the full spectrum of germline AIRR^16^. Besides, gDNA is more stable and easier for storage compared to mRNA, with millions of gDNA samples already stored in the clinical setting or in previous genetic studies. Although mRNA/cDNA-based AIRR can be profiled by many methods, gDNA-based AIRR has just started to attract attention^19–21^.

In this study, we developed the gAIRR Suite comprising three modules (Fig. 1). gAIRR-seq, a probe capture-based targeted gDNA sequencing pipeline, profiles germline AIRR from individual DNA samples. The probe hybridization method can tolerate sequence mismatch to a certain degree and therefore can discover novel alleles. gAIRR-call (Fig. 1), using a coarse-to-fine strategy, starts from aligning gAIRR-seq reads to known AIRR alleles collected from the IMGT database^5^, to eventually call both known and novel AIRR alleles, as well as their flanking sequences. One major challenge for germline AIRR studies is the lack of curated annotations and no ground truth for tool development. We designed gAIRR-annotate to annotate AIRR alleles and flanking sequences, including novel ones, using whole-genome assemblies (Fig. 1). In this study we discovered numerous novel alleles, structural variants, substantial conflicts in different versions of the reference genome, and RSS polymorphism across subjects. The gAIRR Suite can potentially benefit future genetic study and clinical applications for various immune-related phenotypes.

**Fig. 1.**
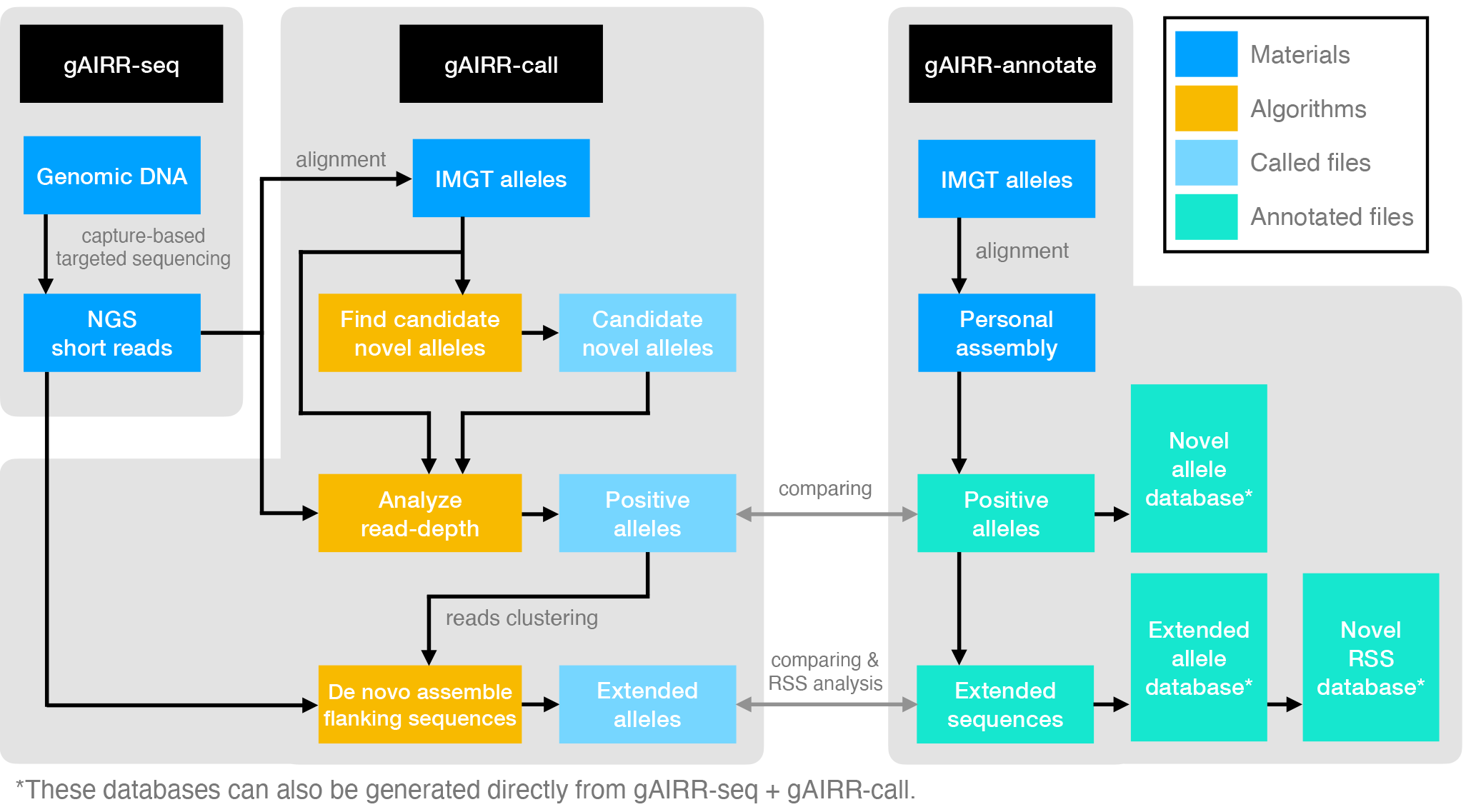
The gAIRR Suite pipelines. The gray arrows show the verification methods between the two pipelines when both gAIRR-seq reads and personal assembly are available. (Methods: Comparing alleles of gAIRR-call and gAIRR-annotate)

## Materials and Methods

### The gAIRR-seq pipeline: a capture-based targeted sequencing method

#### Reference materials

We obtained 7 GIAB genomic DNA RMs from the Coriell Institute (https://www.coriell.org). These 7 RMs included a pilot genome HG001/NA12878 and two Personal Genome Project trios - an Ashkenazim Jewish ancestry: HG002/huAA53E0/ NA24385, HG003/hu6E4515/NA24149, HG004/hu8E87A9/NA24143; a Chinese ancestry: HG005/hu91BD69/NA24631, HG006/huCA017E/NA24694, HG007/hu38168C/NA24695. The genomic DNA samples were retrieved from EBV-transformed B lymphocytes. These RMs were well-sequenced using more than 10 NGS platforms/protocols by GIAB^22^; therefore, they are appropriate RMs for benchmarking gAIRR-seq.

#### Primary cell samples

We also extracted the genomic DNA from both the peripheral blood mononuclear cells (PBMC) and oral mucosal cells of a Taiwanese subject. The subject signed informed consent for participating in the project, and all procedures were approved by the Research Ethics Committee of the National Taiwan University Hospital.

#### Probe design

We designed probes for all known V and J alleles of IG and TR based on the IMGT database Version 3.1.22 (3 April 2019) (including all functional, ORF, and pseudogenes)^5^. Each probe was a continuous 60-bp oligo (Roche NimbleGen, Madison, WI, U.S.). We designed three probes for each V allele and one probe for each J allele based on V and J alleles’ length differences. (Supplementary Fig. S12). For J alleles shorter than 60 bp, we padded the probes with random nucleotides ‘N’ to 60 bp in length on both ends. An example of probes and captured reads’ relative position is shown in Supplementary Section Probe design and captured-reads representation.

#### Library preparation

We quantified the input of 1000 ng gDNA and assessed the gDNA quality using a Qubit 2.0 fluorometer (Thermo Fisher Scientific, Waltham, MA, U.S.) before the library preparation. The gDNA was fragmented using Covaris (Coravis, Woburn, MA, U.S.), aiming at the peak length of 800 bp, which was assessed using Agilent Bioanalyzer 2100 (Agilent Technologies, Santa Clara, CA, U.S.). We then added adapters and barcodes onto the ends of the fragmented DNA to generate indexed sequencing libraries using TruSeq Library Preparation Kit (Illumina, San Diego, CA, U.S.). We performed capture-based target enrichment using the SeqCap EZ Hybridization and Wash Kit (Roche NimbleGen, Madison, WI, U.S.). We then used the sequencing platform, MiSeq (Illumina, San Diego, CA, U.S.), to generate paired-end reads of 300 nucleotides.

### The gAIRR-call pipeline

The gAIRR-call pipeline is composed of three steps: Finding novel alleles, calling alleles, and generating extended alleles with 200 bp flanking sequences on both ends.

#### Finding novel allele candidates

gAIRR-call first generated candidate novel alleles using capture-based short reads. The capture-based short reads were aligned to all the known alleles in the IMGT database (v3.1.22 in this study). Then gAIRR-call checked the reference IMGT alleles one by one. For any known allele, the variants were called with the aligned reads. gAIRR-call marked any allele position with more than a quarter of reads supporting bases different from the reference. If there was only one variant in the reference, gAIRR-call simply called the allele with the variant as a novel allele candidate. If there is more than one variant in the reference, gAIRR-call performs haplotyping and connects the variants with aligned reads. Since most of the AIRR alleles were less than 300 bp in length, there were usually a sufficient number of captured reads (2 × 300 bp in length using gAIRR-seq) for haplotyping.

gAIRR-call then collected called haplotypes and checked if there were duplicated sequences. If a called haplotype was actually another known allele in IMGT, the haplotype was discarded. Because haplotypes were called from known alleles with different lengths, there could be duplicated haplotypes representing the same allele. In this case, only the shorter haplotype was kept. After de-duplicating and cleaning, the remaining haplotypes were unique. The haplotypes joined the known IMGT alleles to be the candidate alleles for the next stage analysis.

#### Calling AIRR alleles

Compared to the previous step, calling AIRR allele applied stricter criteria. gAIRR-call aligned capture-based short reads to the allele candidates pool, which included both known IMGT alleles and the novel ones. To ensure that every candidate allele could be aligned by all potential matched reads, we used the ‘-a’ option of BWA MEM^23^ so that one read could be aligned to multiple positions.

After alignment, gAIRR-call removed alignment that contained mismatches or indels. Further, for alleles longer than 100 bp, the reads’ alignment length should be more than 100 bp. For alleles shorter than 100 bp, the reads should cover the entire allele. Subsequently, only reads perfectly matched the allele with long coverage length passed the filter. gAIRR-call counted the remaining reads’ read-depth of all allele positions. The minimum read-depth in an allele was its final score relative to other candidate alleles. The candidate alleles were sorted according to their supporting minimum read-depth (Fig. 2b).

**Fig. 2.**
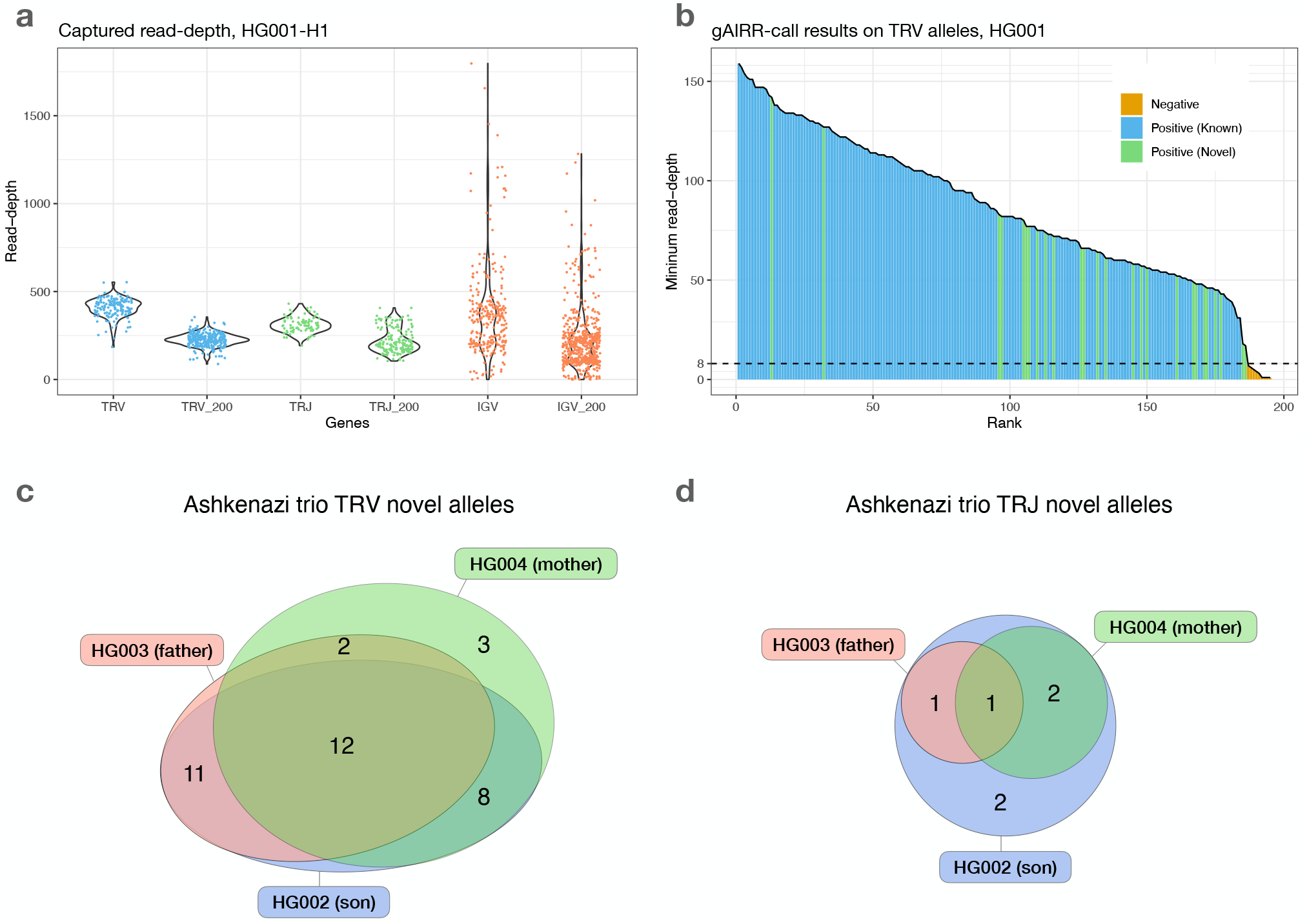
gAIRR-seq and gAIRR-call results. **a**, Read depths sequenced with gAIRR-seq in TRV, TRJ and IGV regions using data from HG001. Columns without the “_200” suffix shows the average read-depth of a region. Columns with the “_200” suffix shows the read depth 200 bp away from the region boundaries. **b**, gAIRR-call results using HG001 data. The results are sorted by minimum read-depth of the perfect matched reads. The dash line represents the adaptive threshold in gAIRR-call. All the alleles not annotated by gAIRR-annotate, colored in orange, are below the adaptive threshold and are regarded as nocalls by gAIRR-call. True known alleles are in blue and true novel alleles are in green. In the HG001 analysis all true alleles are successfully identified by gAIRR-call. Alleles with zero minimum read-depth are not included in the figure. **c**, The TRV and **d**, TRJ novel allele relationship in the Ashkenazi family. HG002: son; HG003: father; HG004: mother. In TRV, there are no unique TRV alleles owned by HG002. In other words, all the HG002’s novel TRV alleles are also owned by either HG003 or HG004. In TRJ, HG002 owns two unique TRJ alleles, which violates Mendelian inheritance. These two alleles are related to *de novo* structural variants (See Section Results: gAIRR-annotate identified in HG002 two structural variants which were further confirmed with gAIRR-seq)

We used an adaptive threshold in gAIRR-call to decide the final calls. The goal of this adaptive threshold was to identify the large drop in minimum read-depth, which was observed to be a good indicator of true-positive and false-positive alleles. When two consecutive alleles had a read depth ratio of less than 0.75, we set the adaptive threshold to be the lower read depth. All the candidate alleles with lower minimum read-depth below the adaptive threshold were discarded. We also noticed that there could be some alleles with extraordinary allele lengths. For example, *TRAV8-5* is a 1355 bp pseudogene that is the longest known TR alleles, where the second-longest allele is 352 bp. When they truly appeared in the data, their minimum read-depths were usually slightly lower than other true alleles’. Thus, when calculating the adaptive threshold, we excluded outlier alleles like *TRAV8-5* to provide a more appropriate threshold. After that, we still compared the minimum read-depth of these outlier alleles with the adaptive threshold to decide whether they were positive.

#### Calling flanking sequences

After allele calling, we performed assembly and haplotyping to call the flanking sequences of the called allele. For each called allele, we grouped the reads perfectly matched with long coverage length with their mate pairs. These reads were then assembled using SPAdes (v3.11.1)^24^ with ‘–only-assembler’ option. We double-check if the allele was actually in the contig by aligning the specific read back to its assembled contigs. The contigs not containing the specific allele were discarded. Mostly, one called allele generated one contig. If multiple contigs were generated, we kept the longest one.

The flanking sequences of an allele could be different in H1 and H2 of a sample. To get the correct phased flanking sequences, we perform haplotyping on the assembled contig. We first aligned all the capture-based short reads to all the contigs. This time, a read could only be aligned to one position. Here we assumed that the flanking sequences of an allele in the two haplotypes were similar and thus the reads of an allele’s two flanking sequences could be aligned to the same contig. We marked the contig position with more than a quarter of reads supporting bases different from the contig. Then we haplotyped the variants using the pair-end reads. The fragment lengths of the pair-end reads were up to 800 bp; hence the core allele region and extended 200 bp from both sides can be fully covered by the pair-end reads. After haplotyping, the extended alleles with 200 bp flanking sequences were reported.

Although the contigs assembled by SPAdes were typically longer than 1000 bp, we did not report the full-length flanking sequences for two reasons. First, the contigs’ boundary regions were not very reliable because the read depth was decreasing toward the boundaries. Second, the fragment length of our capture-based reads was up to 800 bp. If the shortest distance between variants on two flanking sides of the allele is longer than 800 bp, it would not be possible to haplotype the variants. Considering most AIRR alleles are shorter than 300 bp, it is robust to call the extended alleles with 200 bp flanking sequences from both sides.

We performed flanking sequence calling after completion of novel alleles calling for the following reason. If the reads can only be aligned to contigs generated by known alleles, the reads of novel alleles will be aligned to the closest known alleles, resulting in false-positive read assignment and lower accuracy.

### The gAIRR-annotate pipeline

The gAIRR-annotate pipeline annotated AIRR core alleles as well as the extended flanking sequences using publicly available whole-genome assemblies (Supplementary Table S2). gAIRR-annotate begined with aligning IMGT AIRR alleles to the assemblies. To take novel alleles into consideration, we set the alignment options to allow mismatches and indels, and reported all potential sites. For any allele aligned with mismatches or indels, we identified its nearest allele in the IMGT database and assigned a novel allele to the associated gene. When there were multiple assemblies available for the same sample, we only annotated an allele if more than half of the assemblies reported the same output.

To find RSS of each allele, we aligned all IMGT RSS to the annotated flanking sequences. For an RSS aligned with mismatch(es), we defined the aligned region as a novel RSS. The flanking sequences unable to be aligned with IMGT RSS were re-aligned where IMGT heptamers and nonamers were separated. The second-pass processing thus allowed heptamer-nonamer pairs not recorded in IMGT (Supplementary Fig. S15).

For TRV and TRJ alleles, gAIRR-annotate used the core allelic sequences as the query. For TRD alleles, because the lengths of them range from 8 to 16 bp, which would result in a huge number of valid alignments, we extended them by adding the heptamers on both sides of the D region. After the extension with the heptamers, the extended D alleles’ length ranged from 21 to 31. They usually appeared only once in the whole-genome, suggesting the uniqueness of TRD alleles with appropriate RSS.

### HQ-12 set

The HQ-12 set was selected where each sample could either be validated with family information or was curated with additional effort. This dataset includes HG001 (also known as NA12878, Northwest European American from Utah), HG002 (also known as NA24385, Ashkenazi Jewish), Puerto Rican trio (HG00731, HG00732, and HG00733), Southern Han Chinese trio (HG00512, HG00513, and HG00514), Yoruban trio (NA19238, NA19239, and NA19240), and a haploid hydatidiform mole CHM13. All samples have multiple assemblies generated from different technologies, assembly software or research groups. The alleles called using different assemblies from a single sample were concordant in most cases. The pipeline used a conservative strategy for high-precision annotation at conflicted loci – it took the alleles called from more than a half of the assemblies at a locus when the assemblies of a single sample generated inconsistent results. For example, we included six HG002 assemblies in the analysis, so gAIRR-annotate took an allele when there were calls from at least four assemblies. (Supplementary Table S3)

### Comparing alleles of gAIRR-call and gAIRR-annotate

Because both the capture-based sequence data and personal assembly were available in HG001 and HG002, we could compare the results of gAIRR-call and gAIRR-annotate in these two samples. The known alleles identified by gAIRR-call could be directly compared with gAIRR-annotate results; however, comparing the novel alleles were not as straightforward as comparing known ones. As a result, we verified the gAIRR-call novel alleles by aligning them to the personal assemblies. If a novel allele could perfectly align to the assembly, then we considered it as a true positive. We also checked if gAIRR-call alleles covered all the annotated novel allele positions in gAIRR-annotate. Similarly, since there were novel alleles in extended alleles, we evaluated the flanking sequences with alignment rather than direct comparisons. If an annotation is not called by gAIRR-call, it’s regarded as false-negative; if a gAIRR-call result doesn’t have a matched annotation, it’s considered to be false-positive.

### Verifying HG002’s deletion with gAIRR-seq reads

We aligned capture-based short reads of gAIRR-seq from HG002. We first examined if there were reads aligned across two deletion breakpoints, either in a chimeric form or having two paired segments separated across the region. We found 72 reads aligned across chr14:22,918,113 and chr14:22,982,924 using the gAIRR-seq dataset, providing clear evidence of a long deletion.

We also compared the alignment result of HG002 to those of HG003 (HG002’s father) and HG004 (HG002’s mother). We compared the read depth inside and outside the called deletion region. We counted the number of reads in the centromeric and the telomeric regions of chr14:22,982,924 and calculated the coverage drop-off rate (#centromeric reads/#telomeric reads). In a region where reads are nearly uniformly aligned, the drop-off rate is expected to be around 1. If one of the haplotypes is missing on the centromeric side, we expect a drop-off rate around 0.5. At locus chr14:22,982,924, the drop-off rate of HG002 is 0.622, while the values for HG003 and HG004 are 1.005 and 1.138 respectively (Supplementary Table S5). This verified the long deletion in the HG002 TRA/TRD region and further provided evidence that the structural variant was likely a novel mutation.

### Database collecting

We deposited the novel alleles and the extended alleles with 200 bp flanking sequences called by gAIRR-annotate into two databases. Usually, the alleles called by gAIRR-annotate using different assemblies from a single sample are concordant. When there are conflicts between different assemblies, the pipeline uses a conservative strategy for high precision– it takes the alleles called from more than a half of the assemblies at a locus when the assemblies of a single sample generated inconsistent results. For example, we included five HG002 assemblies in the analysis, so gAIRR-annotate took an allele when there were calls from at least three assemblies. We also collected novel alleles and extended alleles called by gAIRR-call. Whenever there are multiple forms of the same allele called, which are inferred from different ways, we keep only the shortest one.

## Results

### gAIRR-seq achieved efficient targeted sequencing in the AIRR regions

We gAIRR-seqed GIAB reference materials (RMs) HG001-7 (EBV-transformed B lymphocytes) and two primary cell samples (PBMC and oral mucosal cells from the same individual) to generate paired-end reads (2 × 300 bp), with a mean of 310 thousand reads for each DNA sample. To evaluate the sequencing quality and efficiency, we aligned the gAIRR-seq reads of the GIAB RMs, HG001 and HG002, to their individual whole-genome assemblies^25^, with over 96% of the reads successfully aligned. Over 83% of the sequenced reads were on-target in the AIRR regions in HG001 and HG002 (Supplementary Table S1). In core TRV and TRJ allelic regions where probes were designed, the average read depth was above 350; the sequencing depth at positions 200 bp away from the allele boundaries was typically above 200x (Fig. 2a). Much higher read depth variation was found in IGV alleles, which might be biased because the DNA was from EBV-transformed B lymphocytes (see Discussion). High on-target rate and high coverage make gAIRR-seq efficient to enrich AIRR sequences and provide high-quality data for computational analyses.

### gAIRR-call could profile known and novel germline TRV and TRJ alleles

We applied gAIRR-call using the high-coverage targeted reads from gAIRR-seq. We collected known alleles in the IMGT database and used them to identify candidate alleles. We define an allele call to be positive if the candidate allele has higher read coverage than a dataset-dependent adaptive threshold (Section Methods: The gAIRR-call pipeline).

To validate the accuracy of the gAIRR-call results, we compared them with the high-confidence annotations from gAIRR-annotate on HG001 and HG002 using their high-quality whole-genome assemblies (Section Results: Annotating TR alleles using high-quality assemblies). According to the comparison, gAIRR-call called TRV and TRJ alleles with 100% accuracy, including both known and novel alleles, for both HG001 and HG002 (Fig. 2b). Furthermore, we provided additional validation using trio analysis, including an Ashkenazi family (son: HG002, father: HG003 and mother: HG004) and a Chinese family (son: HG005, father: HG006 and mother: HG007). We performed gAIRR-call, including assigning novel alleles, independently on each of the samples of the two trios. We showed that there was no Mendelian violation in either family except two calls from HG002 (at *TRAJ29* and *TRBJ2-3*) (Fig. 2c,d and Supplementary Fig. S1). We inspected the violated alleles and noticed they both located near known structural variant breakpoints, which were both regarded as *de novo* mutations (see Section Results: gAIRR-annotate identified in HG002 two structural variants which were further confirmed with gAIRR-seq for further discussion). In a similar fashing, we compared the gAIRR-call results between PBMCs and mucosal cells from the same individual. Out of 174 TRV alleles, 86 TRJ alleles and 22 IGJ alleles, the calling results from PBMCs and mucosal cells are all the same with both known and novel alleles except one TRV allele. In the mucosa calling result, an additional TRV allele *TRDV2*01* was considered positive just above the adaptive threshold. In 396 known IGV alleles, there are only 4 disagreement between PBMCs and mucosal cells. Thus, we conclude that the combined gAIRR-seq and gAIRR-call pipeline is accurate in both known and novel alleles.

Further, we used gAIRR-call to analyze TRV alleles’ flanking sequences outside the core regions, which are important for the V(D)J recombination mechanism but have not been comprehensively collected in the database. Similar to verifying core alleles, we utilized gAIRR-annotate to provide truth sets from whole-genome assemblies. Among the 224 alleles called with flanking sequences from HG001, there were 7 (3%) false positives and 0 false negatives; among the 222 alleles called from HG002, there were only 2 (1%) false positives and 0 false negatives. We further checked if the generated flanking sequences carried appropriate RSS, which were critical in V(D)J recombination. All 7 GIAB RMs and the primary cell sample carried identical or nearly identical RSS with known patterns in IMGT.

It is worth mentioning that we considered three gAIRR-seqed/gAIRR-called extended alleles correct though they did not align perfectly to the personal assemblies HG001 and HG002. We found that, after manual inspection, in all three instances the personal assemblies contained indel errors because of PacBio sequencing weakness in homopolymers (Supplementary Section The flanking sequences where gAIRR-seq and gAIRR-call disagree personal assembly).

### gAIRR-annotate could profile AIRR alleles using genome assemblies

We collected 106 haplotypes from 6 works of literature^25–30^ and annotated TR alleles using gAIRR-annotate. The dataset includes 36 individuals with a broad population background diversity. We first selected 12 extremely high-quality samples to form a “HQ-12” set, where each sample could either be validated with family information or was curated with additional effort (See Section Methods: HQ-12 set). We showed that all haploids had 139-158 TRV alleles, and all but one haploid from HG002 had 5 TRD alleles and 82-86 TRJ alleles (Supplementary Table S2). gAIRR-annotate demonstrated HG002 to have fewer TR alleles, which is consistent with the known two long deletions in the paternal haplotype in the TRD and TRJ region^31^ (See Section Result: gAIRR-annotate identified in HG002 two structural variants which were further confirmed with gAIRR-seq for details). We then gAIRR-annotated the remaining 24 samples from HGSVC^26^ and showed that the allele numbers were similar (Supplementary Table S4).

The novel alleles discovered by gAIRR-annotate showed concordance with the gene locations of existing IMGT annotations. For example, we found a novel allele between *TRAJ15*01* and *TRAJ17*01* in HG001-H1. The novel allele had only one base difference compared to *TRAJ16*01*, which resides in the middle of *TRAJ15* and *TRAJ17* according to IMGT. Thus, we were confident that the allele was a novel allele of a known gene. On average, there were 18 novel V alleles and 3 novel J alleles in each haplotype (Supplementary Table S2). We examined the alleles’ relative positions for HG001 (Fig. 3a) and HG002, and the TRA and TRD alleles’ relative positions were in the same pattern as the *Locus representation of IMGT*^32^ (Supplementary Fig. S3). It shows that the novel alleles found by gAIRR-annotate, marked by purple text, are known genes with a limited number of genetic alterations, mostly single nucleotide substitutions, to the known allele cataloged in IMGT. We also verified the structural difference found by gAIRR-annotate by comparing it with IMGT. For example, both known structurally different combinations of TRGV alleles (12 and 14 genes) in IMGT were found in HG001’s haplotypes. The variations were in concordance with the IMGT gene locus map (Supplementary Fig. S4).

**Fig. 3.**
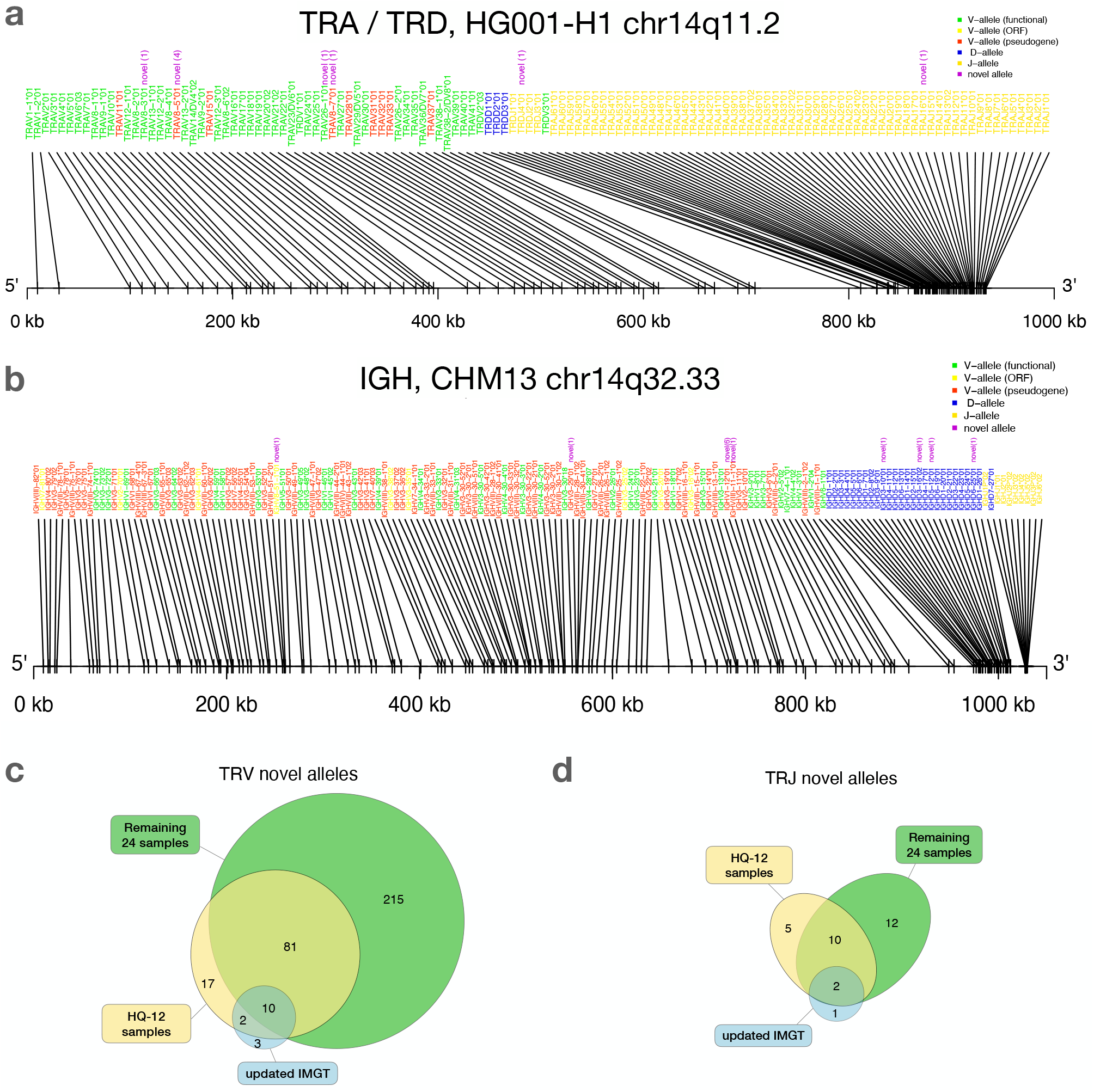
gAIRR-annotate results. **a**, Positions of HG001’s TRA and TRD alleles determined by gAIRR-annotate. The purple text indicates that the allele is novel with its edit-distance compared to the most similar base allele inside parentheses. The text color of the gene names are basically according to IMGT’s color menu for genes^5^. **b**, Positions of CHM13^30^’s IGH alleles determined by gAIRR-annotated. The figure settings are the same as in **a. c**, TRV and **d**, TRJ novel alleles found in HQ-12 samples (shown in yellow) and remaining 24 samples (shown in green) compared to the novel alleles updated in IMGT from v3.1.22 to v3.1.33.

For IG alleles, after avoiding genome assemblies from long-term cultured EBV transformed B lymphocytes, we only applied gAIRR-annotate to the CHM13 genome. We identified 32 novel alleles from the IGH, IGK, and IGL locus. In addition, there are also 32 novel alleles from the orphons of CHM13 (Fig. 3b; Supplementary Table S6).

### Identified novel alleles are consistent with, and far outnumbering, the latest IMGT update

We identified a large number of novel alleles from the 36 samples, as compared with the IMGT database. We used IMGT v3.1.22 (April, 2019) as the baseline in this study, since it was the database we used to design gAIRR-seq probes. The latest version of the IMGT database (v3.1.33, March, 2021) had 15 TRV and 3 TRJ alleles added for human TR since v3.1.22. The novel alleles identified by gAIRR-annotate far outnumbered the IMGT updates (325 vs. 15 for TRV and 29 vs. 3 for TRJ; Fig. 3c,d). For the alleles updated by IMGT, a large fraction of TRV and TRJ alleles (14 out of 18) was also discovered by gAIRR-annotate (Fig. 3c,d), demonstrating high efficiency and accuracy of gAIRR Suite.

The gAIRR-annotate method can also annotate extended TR alleles, which we defined as the core alleles plus 200-bp flanking sequences from both ends. We annotated 511 extended TRV alleles in the HQ-12 set, all of which were phased, and only 413 were known in the latest IMGT database. We further analyzed the RSS in the extended regions given their important participation in the V(D)J recombination mechanism. According to IMGT gene-DB, most TR RSSs follow the rule that the heptamer and nonamer in the RSS are separated by either a 23-bp (191 V alleles) or a 12-bp spacer (89 J alleles). However, there are some functional TR alleles having 24-bp (10 V alleles) or 22-bp (4 V alleles) spacers and occasionally 11-bp (*TRAG20*01*), 10-bp (*TRAJ7*01*) or 13-bp (*TRDJ2*01*) spacers. We used gAIRR-annotate to identify known RSS patterns in the 36 genomes and discovered 56 novel TRV and 20 novel TRJ RSSs (Table 1). We observed RSS polymorphism across different populations. For example, all 36 samples carry the IMGT *TRAJ12*01* RSS and 6 out of 36 samples also carry a novel *TRAJ12*01* RSS with a SNP at the nonamer (heterozygous).

**Table 1.** Number of known and novel TR RSS in the gAIRR-called and gAIRR-annotated flanking sequences. The number of RSS known in IMGT^5^ and novel RSS are shown in columns *#known* and *#novel* respectively. The functionality *Func* indicates that the RSSs come from functional genes while *P/O* indicates that the RSSs come from pseudogenes or open reading frames (ORFs). The RSS from HG001-7 and the primary cell sample are called from both gAIRR-seq and gAIRR-call while the RSS from HQ-12 and the additional 24 samples are called from gAIRR-annotate alone.

| sample | TRV |  |  |  | TRJ |  |  |  |
| --- | --- | --- | --- | --- | --- | --- | --- | --- |
|  | #known |  | #novel |  | #known |  | #novel |  |
|  | Func | P/O | Func | P/O | Func | P/O | Func | P/O |
| HG001 | 94 | 43 | 4 | 1 | 66 | 10 | 7 | 1 |
| HG002 | 95 | 43 | 6 | 1 | 65 | 10 | 9 | 1 |
| HG003 | 94 | 44 | 3 | 1 | 66 | 10 | 6 | 1 |
| HG004 | 91 | 43 | 7 | 1 | 65 | 10 | 8 | 1 |
| HG005 | 93 | 44 | 8 | 2 | 67 | 10 | 7 | 1 |
| HG006 | 96 | 46 | 6 | 1 | 67 | 10 | 8 | 1 |
| HG007 | 95 | 44 | 7 | 2 | 67 | 10 | 5 | 1 |
| Primary cell sample | 95 | 43 | 6 | 2 | 67 | 10 | 6 | 1 |
| HQ-12 set samples | 96 | 46 | 14 | 3 | 67 | 10 | 11 | 3 |
| HGSVC-additional-24 samples | 96 | 46 | 31 | 22* | 67 | 10 | 15 | 3 |

\*: In addition to 22 pseudogene and ORF alleles with novel RSS, we didn't find any appropriate nonamers at *TRBV24/OR9-2\*03* (ORF) for HG01505.

We built a novel-allele database collecting all identified novel alleles, an extended-allele database collecting alleles with 200-bp flanking sequences, and an RSS database collecting RSS information in flanking sequences for both TRV and TRJ (novel-allele: Supplementary File S1 to S4; extended-allele: Supplementary File S5 to S8; RSS: Supplementary File S9 to S12). We showed that we could broadly expand the size of known TR databases using gAIRR-annotate and high-quality whole-genome assemblies. With the rapidly increasing number of high-quality assemblies utilizing accurate long-read sequencing, we are optimistic that gAIRR-annotate will help the community uncover TR germline DNA information at a much greater speed than traditional methods.

### gAIRR-annotate identified in HG002 two structural variants which were further confirmed with gAIRR-seq

From Supplementary Table S2, we noticed low numbers of TRJ and TRD alleles in the paternal haplotype of HG002. When visualizing alleles identified by gAIRR-annotate, we observed a 65 kbp missing region in TRA/TRD (Fig. 4a,b,c), containing 1 D, 1 V, and 36 J alleles from *TRDD3* to *TRAJ30* in the paternal haplotype. Similarly, we noticed a 10 kbp missing TRB region, containing 2 D and 9 J alleles from *TRBD1* to *TRBJ2-2P*, in the same haplotype (Supplementary Fig. S5b).

**Fig. 4.**
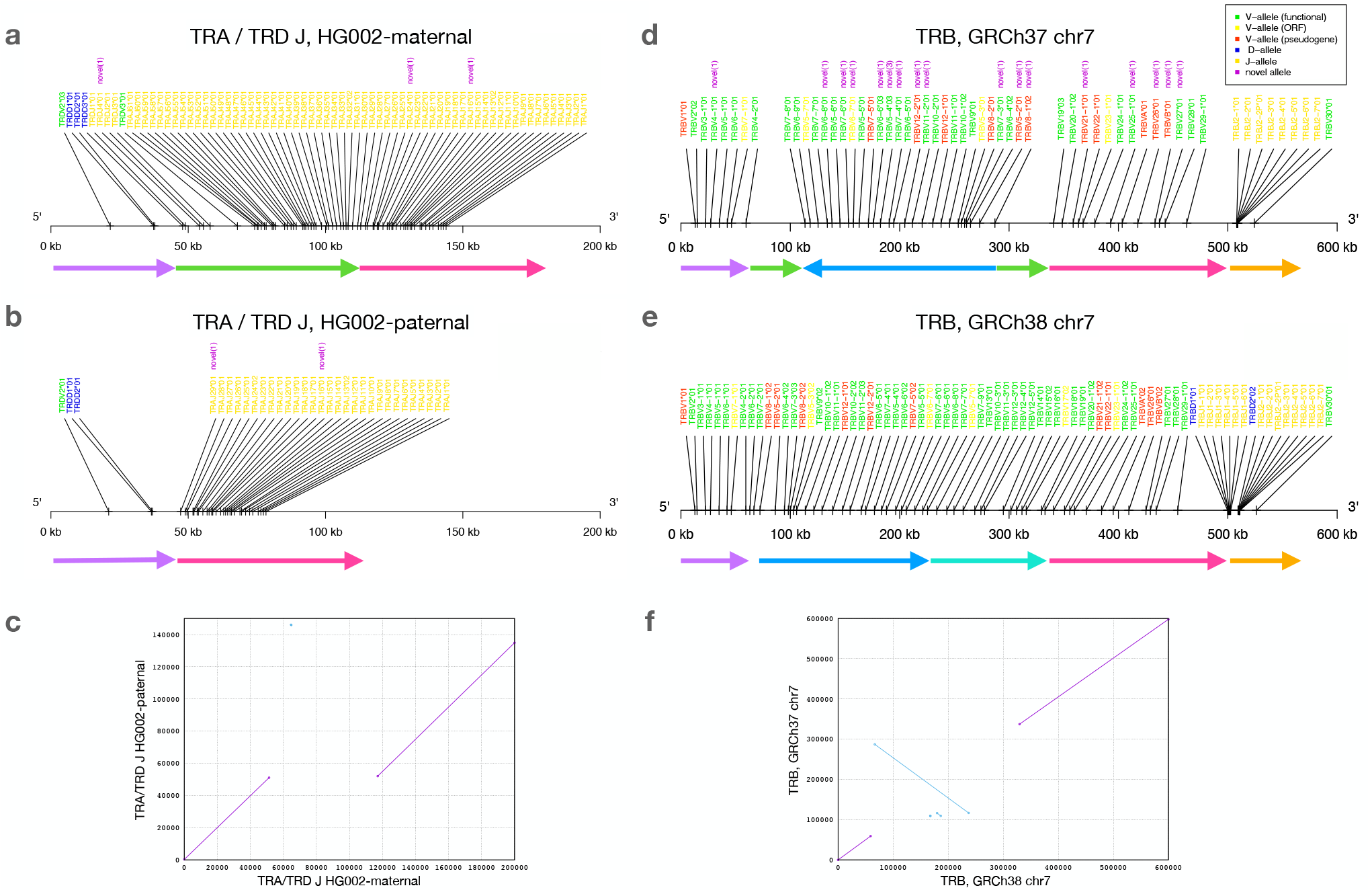
Structural variants found by gAIRR-annotate. **a**,**b**,**c**, The 65 kbp structural variation in TRA and TRD J region of HG002. **a**: maternal haplotype, **b**: paternal haplotype, and **c**: The sequence alignment of HG002’s maternal and paternal TRA/TRD J region. In **a** and **b**, the arrows with the same color can be aligned between the two haplotypes. The green arrow is the segment deleted in paternal haplotype. **d**,**e**,**f**, The inversion and deletion of TR beta chain germline genes of the reference genome. **d**: GRCh37 chr7, **e**: GRCh38 chr7, and **e**: The sequence alignment of GRCh37 and GRCh38 at TR beta chain. In **d** and **e**, the arrows with the same color can be aligned between the two haplotypes. The deep blue arrow indicates the inversion between the reference genomes.

To further verify the 65 kbp deletion in the TRA/TRD loci of HG002, we aligned capture-based short reads of gAIRR-seq from HG002, as well as gAIRR-seq reads from HG003 (father) and HG004 (mother), to GRCh37 for a trio analysis (Fig. 5). By examining the split reads across a certain region, and also the read depth reduction within a continuous region, we confirmed the *de novo* deletion in HG002 (see Section Methods: Verify HG002’s deletion with gAIRR-seq reads). The deletion contributes to a loss of 32 TRAJ genes and all 4 TRDJ genes from the paternal haplotype.

**Fig. 5.**
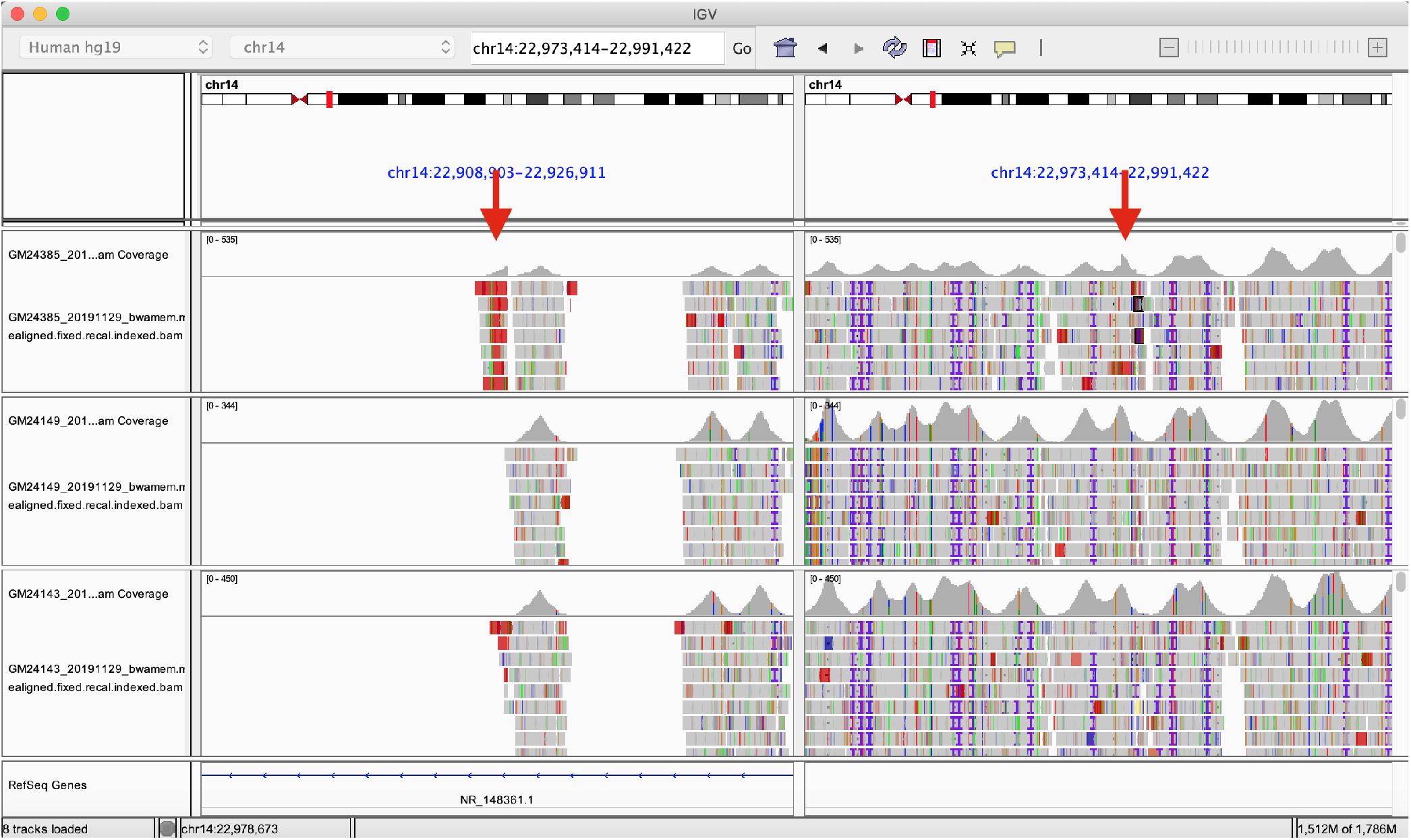
The Integrated Genomics Viewer visualization of HG002’s structural variant. The Integrated Genomics Viewer visualization of the capture-based reads from HG002 (son, top), HG003 (father, middle), and HG004 (mother, bottom) aligned to GRCh37 chromosome 14. There are two red arrows indicating abrupt read-depth changes of HG002’s reads at chr14:22,918,113 and chr14:22,982,924.

Our results indicated a long deletion on chr14 from position 22,918,114 to position 22,982,924 (GRCh37) for one HG002 haplotype. The structure variant analysis performed by GIAB^31^ also showed a deletion at chr14:22,918,114 in GRCh37, where a 13 bp fragment in HG002 replaces a 64,807 bp fragment in GRCh37.

Similarly, there is a 10 kb deletion on chromosome 7q34 related to TRB alleles. We verified the deletion with gAIRR-annotate, gAIRR-seq, and the report from GIAB^31^. (Supplementary Section HG002’s deletion on chromosome 7)

### gAIRR-annotate uncovered substantial conflicts in GRCh37 and GRCh38

We identified an inversion in GRCh37 with respect to the GRCh38 primary assembly that covered TRBV alleles (Fig. 4d,e,f). In this inversion, there were 39 TRBV genes (*TRBV6-2* to *TRBV18* following the gene order provided by IMGT) successfully annotated in GRCh38. In contrast, in GRCh37, only 27 genes from *TRBV8-1* to *TRBV7-8* were successfully annotated, but with a reversed order. We observed another long missing sequence (about 10 kbp) in GRCh37 in the TRB region, covering about half of the expected TRBD and TRBJ genes. We noticed GRCh38 didn’t carry this deletion, and we annotated reasonable numbers of TRBD and TRBJ genes. When comparing annotated alleles to the IMGT database, we showed that on chromosome 7, all alleles in GRCh38 are known, and there are 18 novel alleles in GRCh37. In IG alleles, there are also some differences between the reference genomes; however, all the genome structures are in consistence with IMGT’s locus representation. (Supplementary Fig. S8; Supplementary Table S6)

We further analyzed an GRCh38 alternative contig, *chr7_KI270803v1_alt*, which contained TRBV alleles (Supplementary Fig. S9). We identified eleven novel TRBV alleles in *chr7_KI270803v1_alt* using gAIRR-annotate. Compared to the GRCh38 primary assembly, *chr7_KI270803v1_alt* has six additional TRBV genes. TRB genes in both chr7 and *chr7_KI270803v1_alt* showed concordance with the *locus representation* of IMGT^32^. Thus, when a personal assembly is not available, we suggest using GRCh38 instead of GRCh37 for TR analysis in that GRCh38 shows concordance with the IMGT database in sequence completeness and carries better-known alleles.

## Discussion

We have developed the gAIRR Suite that includes probe capture-based targeted sequencing and computational analyses to profile germline AIRR. gAIRR-seq can enrich the AIRR regions with high on-target rates and sufficient read depth, and gAIRR-call can then accurately call both known and novel alleles using gAIRR-seq reads. gAIRR-annotate can take advantage of whole genome assemblies to accurately annotate AIRR genes in both core allelic and flanking regions. We applied gAIRR-annotate to identify 325 novel core alleles in TRV and 29 novel core alleles in TRJ from 36 subjects. We identified RSSs with spacers neither 12-bp nor 23-bp in length, which had been reported before^33,34^ but not yet comprehensively profiled. gAIRR-annotate can also help to discover structural variants. We independently identified and then verified two known structural variants in the TR regions in HG002. We also searched for structural variants in the IG regions of CHM13, and found that the genome structures are consistent with IMGT’s locus presentation. Similarly, we used gAIRR-annotate to uncover substantial conflicts between references GRCh37 and GRCh38.

gAIRR-seq is advantageous in its high resolution and comprehensiveness. Compared to multiplex PCR-based methods, gAIRR-seq covers all known AIRR genes and alleles in one experiment. Furthermore, novel alleles can also be identified because the probes can tolerate sequence mismatch to a certain degree. Each sample only costs approximately USD 200 for library preparation, target enrichment, and sequencing while all IG and TR genes/alleles being gAIRR-seqed in one experiment using easily available genomic DNA from cell lines or primary cells, such as PBMCs or mucosal cells. We envision that gAIRR-seq capture probe design can be further upgraded by gAIRR-annotate results thanks to the collection of accurate extended alleles. In this study, our probe design was limited to only V and J genes due to insufficient allele lengths for other genes. With a comprehensive population genomic AIRR database becoming available, we plan to include TRD, IGD and IGJ genes in future gAIRR-seq probe design.

Although the gAIRR-seq and gAIRR-call pipeline is technically applicable to IG genes, we found that in this study the accuracy for IG genes was lower. We reason that the seven GIAB DNA samples were retrieved from EBV-transformed B lymphocytes and might suffer from reduced diversity during the establishment, growth and subculture, making them unsuitable for germline IG profiling^35^. To solve this problem, our strategy is to take advantage of primary cells for IG experiments. In this study, we gAIRR-seqed the genomic DNA from both PBMCs and mucosal cells of a Taiwanese subject (Supplementary Section gAIRR-seq and gAIRR-call on primary cells) and gAIRR-called similar numbers of known and novel alleles in the TRV and TRJ regions compared to the GIAB RMs. We reason that, in PBMCs, B lymphocytes and monocytes can provide un-recombined V(D)J genes for TR sequencing, and T lymphocytes and monocytes can provide un-recombiend V(D)J genes for IG sequencing. Since the gAIRR-call result on the PBMCs and mucosal cells show high concordance, utilizing primary cells (either PBMC or mucosal cells) for IG and TR genotyping is valid.

Up to now, it is almost insurmountable to study the impact of germline AIRR variations on human immune-related phenotypes and diseases. The single nucleotide polymorphism (SNP) genotyping array widely used for genome-wide association study (GWAS) tags only a tiny fraction of AIRR variations^3,12,14^. Whole-genome sequencing (WGS) can theoretically cover AIRR variations; however, with neither a powerful bioinformatics tool (such as gAIRR-call) nor reliable reference genome(s)^15^, WGS has not yet been successfully used to retrieve AIRR information for human genetic study due to high genomic complexity in the AIRR regions and relatively lower sequencing depth compared to targeted sequencing. Whether germline AIRR variations have a broad and strong impact on human immune-related phenotypes and diseases, just like the human leukocyte antigen (HLA) variations do^36,37^, is an attractive question awaiting more studies. In our view, today’s status of germline AIRR is similar to the situation of SNP before the International HapMap Project^38^, or HLA before Sanger sequencing-based typing (SBT). We badly need a HapMap equivalent for germline AIRR.

We envision that gAIRR Suite can help in at least the following scenarios. First, the discovered novel alleles can facilitate development of better SNP arrays and better WGS analysis references of AIRR in the future. Second, the personal germline AIRR profile can be directly employed for genetic study and clinical applications. Third, the personal germline AIRR genes, together with other critical immune genes (such as HLA and killer cell immunoglobulin-like receptor (KIR) genes), can be tested for di-genic or oligo-genic effects on various immune-related phenotypes. Last but not least, the combined analysis of germline AIRR and dynamic mRNA-based AIRR profiles might help decipher molecular mechanisms underlying immunological responses.

## Supporting information

Supplemental Section 1 to 16

## Data Availability

The gAIRR-seq sequence data of the seven reference materials and two primary cell samples have been deposit in Sequence Read Archive https://www.ncbi.nlm.nih.gov/sra/PRJNA767687. The source code of gAIRR Suite is publicly available on GitHub at https://github.com/maojanlin/gAIRRsuite under the *scripts/* directory. The supplementary files including the novel allele, flanking sequence, and RSS databases, and the annotation bed files of human reference genome GRCh37 and GRCh38 (Details in Supplementary Section Annotation files of the human reference genomes) are under the directory *supplementary_files/*.

## Acknowledgements

We thank to National Core Facility for Biopharmaceuticals (NCFB, MOST 108-2319-B-492 -001) for support and National Center for High-performance Computing (NCHC) of National Applied Research Laboratories (NARLabs) in Taiwan for providing computational and storage resources. This study was supported by Taiwan Ministry of Science and Technology grants (MOST 108-2314-B-002-069-MY3) and National Taiwan University Hospital (109-S4521, 108-T05). We also thank Justin Zook for his helpful suggestions in sequencing experiments and analysis.

## Author contributions statement

Y.L. and P.C. designed the sequencing experiments and A.C.L. performed the experiments. Y.L. and N.C. designed the preliminary analysis pipeline. M.L. designed the final analysis pipelines and wrote the software. M.L., N.C., and Y.L. interpreted all the analysis results with P.C.’s guidance. S.K., J.H., C.C., C.H., W.Y., and P.C. initiated this project and assisted with discussions. M.L., N.C., and P.C. wrote the manuscript with input from all authors. All authors read and approved the final manuscript.

