## Supplemental Section 1 to 16 for "Profiling Germline Adaptive Immune Receptor Repertoire with gAIRR Suite"

#### Supplemental Materials

### 1 gAIRR-seq's result on 7 RMs and two primary cell samples

**Table S1.** The number of reads sequenced, fraction of aligned reads and on-target rate of gAIRR-seq using samples HG001 and HG002. The target region is defined as the union of annotated allelic regions, where each region includes the core allele and 800 bps extended from both sides as shown in Supplementary Fig. S13. RM HG003-7 and the primary cell sample are not applicable in capture-efficiency analysis for lacking personal assembly annotation.

| Sample | # of reads sequenced | % of reads aligned (H1/H2) | % in target region (H1/H2) |
|---|---|---|---|
| HG001 | 280,579 | 97.6/98.0 | 83.3/83.7 |
| HG002 | 281,494 | 98.5/96.1 | 84.5/83.3 |
| HG003 | 307,312 | N/A | N/A |
| HG004 | 362,271 | N/A | N/A |
| HG005 | 327,354 | N/A | N/A |
| HG006 | 338,361 | N/A | N/A |
| HG007 | 297,186 | N/A | N/A |
| Primary cell sample | 283,691 | N/A | N/A |

We located all TRV, TRJ, and IGV alleles in the assemblies with gAIRR-annotate to validate if the reads are in AIRR regions. Considering the fragment lengths are up to 800 bp in our sequencing method, we defined a read aligned within 800 bp from its allele boundaries to be on-target.

We assessed the biases of probe preferences during library preparation using probe capture-based enrichment. The only bias we observed is an extremely long TRV pseudogene *TRAV8-5*. The gene length is 1355 bp, while the second-longest TRV gene is only 352 bp. The read-depth coverage of *TRAV8-5*, 270x, is relatively low compared to other V alleles, 405x on average. gAIRR-call tackles the uneven problem by making *TRAV8-5* a special case in adaptive threshold generation (Section Methods: The gAIRR-call pipeline). The lower read coverage of *TRAV8-5* is likely due to the high allele length-to-probe ratio. By adding more probes according to the allele length, the bias can probably be solved.

#### 2 The novel TRV and TRJ allele relationship in the Chinese trio

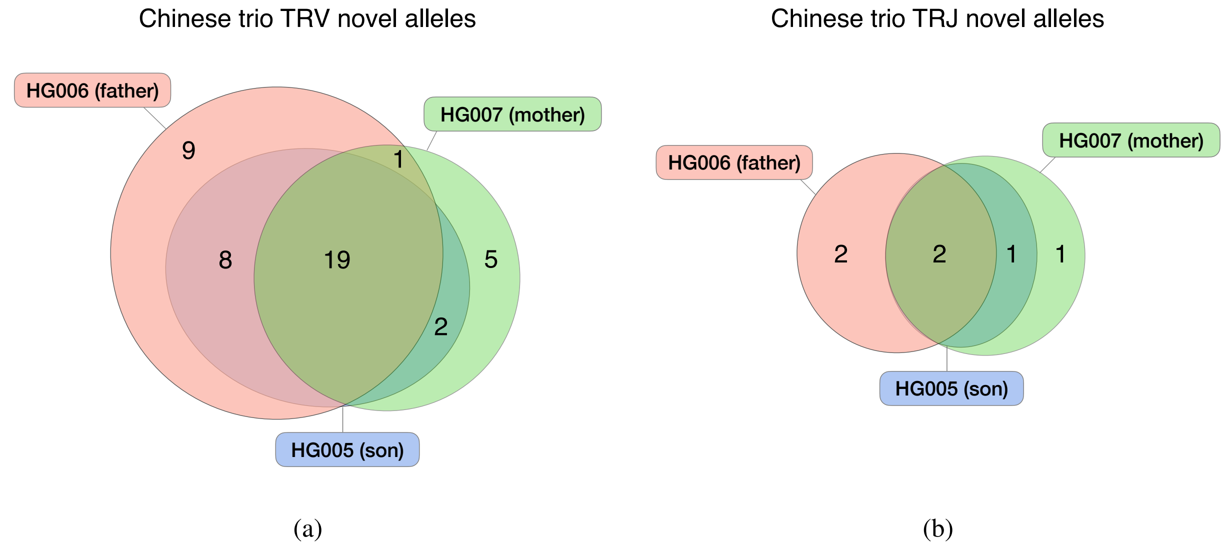


**Fig. S1.** The (a) TRV and (b) TRJ novel allele relationship in the Chinese family. HG005: son; HG006: father; HG007: mother.

#### 3 The flanking sequences where gAIRR-seq and gAIRR-call disagree personal assembly

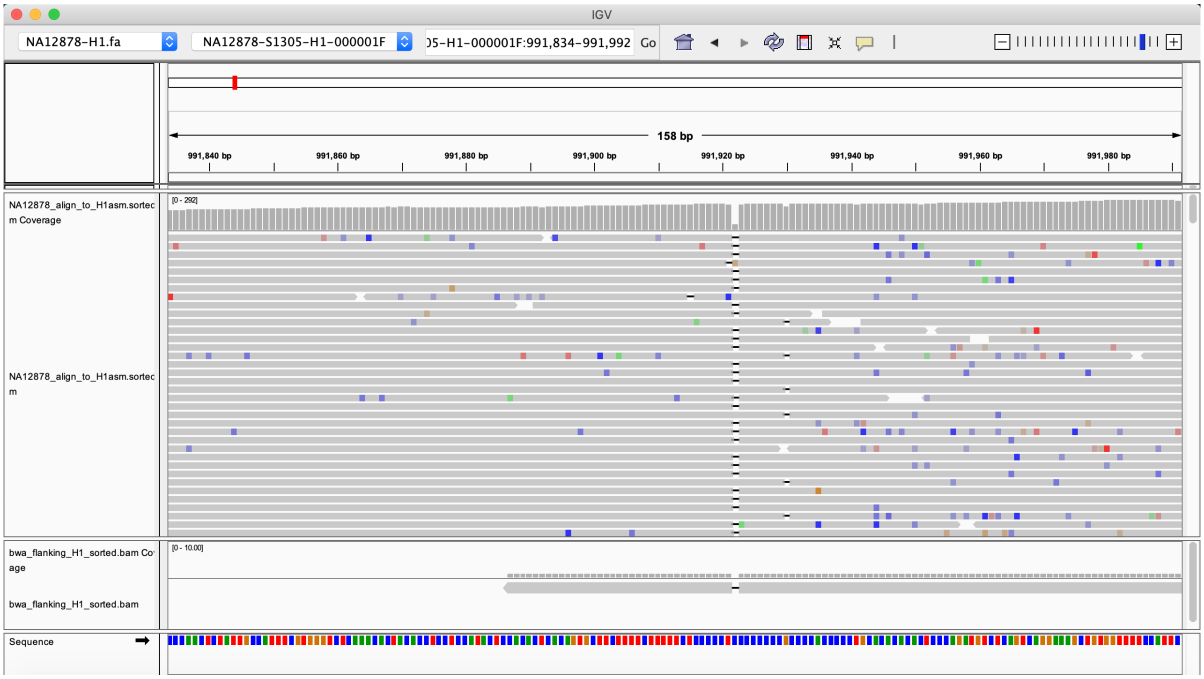


**Fig. S2.** The Integrated Genomics Viewer[39] visualization of the position where gAIRR-seq and gAIRR-call disagree with the HG001 assembly[25]. The reference is the H1 of the assembly. The aligned sequences from top to bottom are gAIRR-seq reads and gAIRR-called flanking sequences aligned to the assembly. The disagreement at position 991922 is homopolymers C where the assembly is 8 consecutive guanines while gAIRR-seq and gAIRR-call support 7 consecutive guanines.

There are three extended alleles not aligned perfectly to the personal assemblies HG001 and HG002, but we still considered them as true positives after manual inspection. The genes *TRBV20-1* in HG001-H1, *TRBV17* in HG002-maternal, and *TRBV20-1* in HG002-paternal all carried one indel with respect to the personal assemblies. All three indels were in homopolymeric regions, which were known to be error-prone using PacBio sequencing. For example, in *TRBV17* the called allele had 7 consecutive guanines in the flanking region, while the assembly had 8 consecutive guanines (Supplementary Fig. S2 We examined these regions with the deep-coverage read sets sequenced with gAIRR-seq using the Integrated Genomics Viewer[39] and concluded the called alleles were likely to be correct.

#### 4 gAIRR-annotate results on collected samples

**Table S2.** Number of TR alleles for HQ-12 set (from 12 samples with 58 haplotypes). The numbers of annotated novel alleles and total alleles are shown in columns *#novel* and *#total* respectively.

| | TRV | | TRD (+hep) | | TRJ | |
|---|---|---|---|---|---|---|
| haplotype | #novel | #total | #novel | #total | #novel | #total |
| HG001-H1[25] | 14 | 145 | 0 | 5 | 2 | 84 |
| HG001-H2[25] | 19 | 146 | 0 | 5 | 3 | 84 |
| HG001-CCS-H1[26] | 20 | 145 | 0 | 5 | 3 | 84 |
| HG001-CCS-H2[26] | 11 | 144 | 0 | 5 | 2 | 84 |
| HG002-P[25] | 23 | 144 | 0 | 2 | 4 | 39 |
| HG002-M[25] | 17 | 147 | 0 | 5 | 3 | 84 |
| HG002-CCS-Canu-P[27] | 24 | 144 | 0 | 2 | 4 | 39 |
| HG002-CCS-Canu-M[27] | 19 | 148 | 0 | 5 | 3 | 84 |
| HG002-CCS-Falcon-P[27] | 20 | 144 | 0 | 2 | 4 | 39 |
| HG002-CCS-Falcon-M[27] | 21 | 146 | 0 | 5 | 4 | 84 |
| HG002-CCS-wtdbg2-P[27] | 23 | 142 | 0 | 2 | 4 | 39 |
| HG002-CCS-wtdbg2-M[27] | 18 | 145 | 0 | 5 | 3 | 82 |
| HG002-hifiasm-P[28] | 23 | 144 | 0 | 2 | 4 | 77 |
| HG002-hifiasm-M[28] | 17 | 147 | 0 | 5 | 3 | 84 |
| HG002-CCS-H1[26] | 21 | 145 | 0 | 2 | 5 | 75 |
| HG002-CCS-H2[26] | 19 | 147 | 0 | 5 | 3 | 84 |
| HG00731-CCS-H1[26] | 12 | 143 | 0 | 5 | 3 | 84 |
| HG00731-CCS-H2[26] | 8 | 144 | 0 | 5 | 2 | 84 |
| HG00731-CLR-H1[26] | 10 | 144 | 0 | 5 | 3 | 84 |
| HG00731-CLR-H2[26] | 12 | 145 | 0 | 5 | 2 | 84 |
| HG00732-CCS-H1[26] | 16 | 147 | 0 | 5 | 1 | 84 |
| HG00732-CCS-H2[26] | 11 | 145 | 0 | 5 | 0 | 84 |
| HG00732-CLR-H1[26] | 16 | 147 | 0 | 5 | 1 | 84 |
| HG00732-CLR-H2[26] | 14 | 146 | 0 | 5 | 0 | 84 |
| HG00733-hifiasm-H1[28] | 13 | 143 | 0 | 5 | 3 | 84 |
| HG00733-hifiasm-H2[28] | 11 | 145 | 0 | 5 | 0 | 84 |
| HG00733-HiFi-v0-H1[29] | 13 | 143 | 0 | 5 | 3 | 84 |
| HG00733-HiFi-v0-H2[29] | 11 | 145 | 0 | 5 | 0 | 84 |
| HG00733-CCS-H1[26] | 11 | 145 | 0 | 5 | 0 | 84 |
| HG00733-CCS-H2[26] | 13 | 143 | 0 | 5 | 3 | 84 |
| HG00733-CLR-H1[26] | 12 | 145 | 0 | 5 | 3 | 84 |
| HG00733-CLR-H2[26] | 14 | 145 | 0 | 5 | 0 | 84 |
| HG00512-CCS-H1[26] | 10 | 141 | 0 | 5 | 3 | 84 |
| HG00512-CCS-H2[26] | 20 | 144 | 0 | 5 | 5 | 84 |
| HG00512-CLR-H1[26] | 16 | 141 | 0 | 5 | 5 | 84 |
| HG00512-CLR-H2[26] | 16 | 144 | 0 | 5 | 3 | 84 |

| | TRV | | TRD (+hep) | | TRJ | |
|---|---|---|---|---|---|---|
| haplotype | #novel | #total | #novel | #total | #novel | #total |
| HG00513-CCS-H1[26] | 11 | 144 | 0 | 5 | 3 | 84 |
| HG00513-CCS-H2[26] | 15 | 145 | 0 | 5 | 3 | 84 |
| HG00513-CLR-H1[26] | 13 | 138 | 0 | 5 | 3 | 84 |
| HG00513-CLR-H2[26] | 16 | 138 | 0 | 5 | 3 | 84 |
| HG00514-CCS-H1[26] | 11 | 149 | 0 | 5 | 3 | 84 |
| HG00514-CCS-H2[26] | 15 | 144 | 0 | 5 | 3 | 84 |
| HG00514-CLR-H1[26] | 17 | 146 | 0 | 5 | 3 | 84 |
| HG00514-CLR-H2[26] | 23 | 144 | 0 | 5 | 4 | 84 |
| NA19238-CCS-H1[26] | 32 | 143 | 0 | 5 | 2 | 84 |
| NA19238-CCS-H2[26] | 24 | 145 | 0 | 5 | 3 | 84 |
| NA19238-CLR-H1[26] | 34 | 156 | 0 | 5 | 2 | 84 |
| NA19238-CLR-H2[26] | 30 | 149 | 0 | 5 | 3 | 84 |
| NA19239-CCS-H1[26] | 24 | 144 | 0 | 5 | 2 | 84 |
| NA19239-CCS-H2[26] | 23 | 144 | 0 | 5 | 4 | 84 |
| NA19239-CLR-H1[26] | 22 | 144 | 0 | 5 | 2 | 84 |
| NA19239-CLR-H2[26] | 25 | 144 | 0 | 5 | 4 | 84 |
| NA19240-CCS-H1[26] | 23 | 144 | 0 | 5 | 2 | 84 |
| NA19240-CCS-H2[26] | 22 | 145 | 0 | 5 | 5 | 84 |
| NA19240-CLR-H1[26] | 24 | 144 | 0 | 5 | 2 | 84 |
| NA19240-CLR-H2[26] | 23 | 153 | 0 | 5 | 5 | 84 |
| CHM13-hifiasm[28] | 11 | 147 | 0 | 5 | 1 | 86 |
| CHM13-T2T[30] | 11 | 147 | 0 | 5 | 1 | 86 |

**Table S3.** Number of TR alleles for HQ-12 sample's consensus. An allele sequence is considered positive only if the allele is called from more than half of the assemblies.

| | TRV | | TRD (+hep) | | TRJ | |
|---|---|---|---|---|---|---|
| sample's consensus | #novel | #total | #novel | #total | #novel | #total |
| HG001 | 29 | 188 | 0 | 5 | 4 | 87 |
| HG002 | 31 | 186 | 0 | 5 | 6 | 89 |
| HG00731 | 17 | 174 | 0 | 5 | 4 | 89 |
| HG00732 | 22 | 173 | 0 | 5 | 1 | 88 |
| HG00733 | 21 | 183 | 0 | 6 | 3 | 81 |
| HG00512 | 26 | 183 | 0 | 6 | 6 | 89 |
| HG00513 | 18 | 156 | 0 | 5 | 5 | 88 |
| HG00514 | 19 | 158 | 0 | 5 | 4 | 88 |
| NA19238 | 47 | 198 | 0 | 5 | 5 | 91 |
| NA19239 | 39 | 190 | 0 | 5 | 5 | 91 |
| NA19240 | 42 | 192 | 0 | 5 | 6 | 92 |
| CHM13 | 11 | 147 | 0 | 5 | 1 | 85 |

**Table S4.** Number of TR alleles for the additional-24 assemblies[26].

| | TRV | | TRD (+hep) | | TRJ | |
|---|---|---|---|---|---|---|
| haplotype | #novel | #total | #novel | #total | #novel | #total |
| HG00096-CLR-H1 | 22 | 144 | 0 | 5 | 2 | 84 |
| HG00096-CLR-H2 | 12 | 143 | 0 | 5 | 1 | 84 |
| HG00171-CLR-H1 | 13 | 144 | 0 | 5 | 2 | 84 |
| HG00171-CLR-H2 | 17 | 147 | 0 | 5 | 2 | 84 |
| HG00864-CLR-H1 | 22 | 150 | 0 | 5 | 2 | 84 |
| HG00864-CLR-H2 | 21 | 150 | 0 | 5 | 4 | 84 |
| HG01114-CLR-H1 | 23 | 147 | 0 | 5 | 0 | 84 |
| HG01114-CLR-H2 | 18 | 150 | 0 | 5 | 1 | 84 |
| HG01505-CLR-H1 | 21 | 144 | 0 | 5 | 2 | 84 |
| HG01505-CLR-H2 | 17 | 142 | 0 | 5 | 1 | 84 |
| HG01596-CLR-H1 | 21 | 145 | 0 | 5 | 3 | 84 |
| HG01596-CLR-H2 | 22 | 151 | 0 | 5 | 1 | 84 |
| HG02011-CLR-H1 | 26 | 144 | 0 | 5 | 2 | 83 |
| HG02011-CLR-H2 | 12 | 144 | 0 | 5 | 1 | 84 |
| HG02492-CLR-H1 | 22 | 144 | 0 | 5 | 1 | 82 |
| HG02492-CLR-H2 | 15 | 152 | 0 | 5 | 3 | 84 |
| HG02587-CLR-H1 | 21 | 145 | 0 | 5 | 2 | 84 |
| HG02587-CLR-H2 | 28 | 142 | 0 | 5 | 3 | 84 |
| HG02818-CCS-H1 | 28 | 145 | 0 | 5 | 1 | 84 |
| HG02818-CCS-H2 | 23 | 145 | 0 | 5 | 3 | 84 |
| HG03009-CLR-H1 | 21 | 150 | 0 | 4 | 1 | 84 |
| HG03009-CLR-H2 | 17 | 144 | 0 | 5 | 1 | 84 |
| HG03065-CLR-H1 | 24 | 147 | 0 | 5 | 2 | 84 |
| HG03065-CLR-H2 | 31 | 147 | 0 | 5 | 3 | 84 |
| HG03125-CCS-H1 | 24 | 144 | 0 | 5 | 2 | 84 |
| HG03125-CCS-H2 | 22 | 147 | 0 | 5 | 3 | 84 |
| HG03371-CLR-H1 | 27 | 145 | 0 | 5 | 4 | 84 |
| HG03371-CLR-H2 | 29 | 142 | 0 | 5 | 1 | 84 |
| HG03486-CCS-H1 | 19 | 145 | 0 | 5 | 5 | 84 |
| HG03486-CCS-H2 | 24 | 145 | 0 | 5 | 3 | 84 |
| HG03683-CLR-H1 | 15 | 144 | 0 | 5 | 1 | 84 |
| HG03683-CLR-H2 | 19 | 144 | 0 | 5 | 2 | 84 |
| HG03732-CLR-H1 | 10 | 144 | 0 | 5 | 3 | 84 |
| HG03732-CLR-H2 | 23 | 144 | 0 | 5 | 2 | 84 |
| NA12329-CLR-H1 | 9 | 145 | 0 | 5 | 1 | 84 |
| NA12329-CLR-H2 | 11 | 144 | 0 | 5 | 2 | 84 |
| NA18534-CLR-H1 | 26 | 144 | 0 | 5 | 3 | 84 |
| NA18534-CLR-H2 | 15 | 144 | 0 | 5 | 3 | 84 |
| NA18939-CLR-H1 | 19 | 144 | 0 | 5 | 3 | 84 |
| NA18939-CLR-H2 | 11 | 144 | 0 | 5 | 5 | 84 |
| NA19650-CLR-H1 | 23 | 147 | 0 | 5 | 1 | 84 |
| NA19650-CLR-H2 | 20 | 141 | 0 | 5 | 3 | 84 |
| NA19983-CLR-H1 | 20 | 145 | 0 | 5 | 2 | 84 |
| NA19983-CLR-H2 | 29 | 144 | 0 | 5 | 2 | 84 |
| NA20509-CLR-H1 | 12 | 146 | 0 | 5 | 2 | 84 |
| NA20509-CLR-H2 | 18 | 142 | 0 | 5 | 0 | 84 |
| NA20847-CLR-H1 | 17 | 144 | 0 | 5 | 0 | 84 |
| NA20847-CLR-H2 | 17 | 141 | 0 | 5 | 2 | 92* |

*: NA20847-CLR-H2 seems to have more TRJ alleles than other haplotypes. In gAIRR-annotate details, the assembly NA20847-CLR-H2 has two contigs covering the same TRGJ gene locus. It is either due to NA20847 having extra TRGJ genes or misassembly in the region.

#### 5 TRA/TRD locus representation and version difference of IMGT

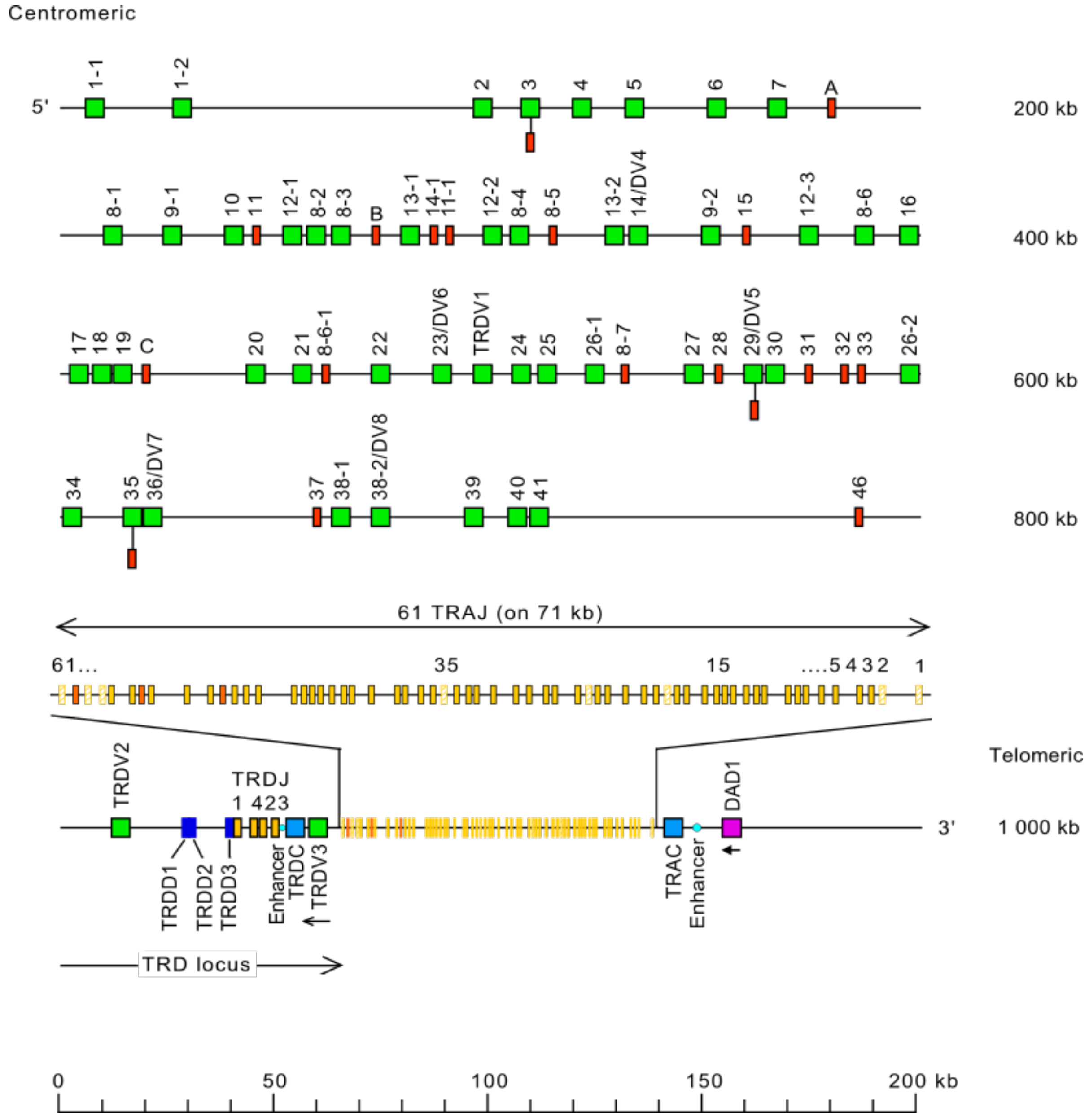


**Fig. S3.** The locus representation of TRA/TRD alleles downloaded from IMGT[32]

There are some pseudogenes, in red, shown in the IMGT locus representation but not found by gAIRR-annotate. It is because the alleles information used by gAIRR-annotate and gAIRR-call is collected from IMGT version 3.1.22 (2019-04-03), while the representation is from the latest IMGT version v3.1.33 (2021-03-22) by the completion of this manuscript. In version 3.1.22, some TR pseudogenes have not been discovered yet. gAIRR-annotate cannot detect genes that are not in the database, so the latest found pseudogenes are missing in the gAIRR-annotate result. In other words, gAIRR-annotate can only detect novel alleles but not novel genes. After the updates from 3.1.22 to 3.1.33, there are 26 more human TRV alleles and three more human TRJ alleles. In the 26 novel alleles updated by the IMGT, 11 are alleles from 8 novel genes (all pseudogenes), which are not considered by gAIRR-annotate.

#### 6 TRG germline variation of HG001 concordant with IMGT

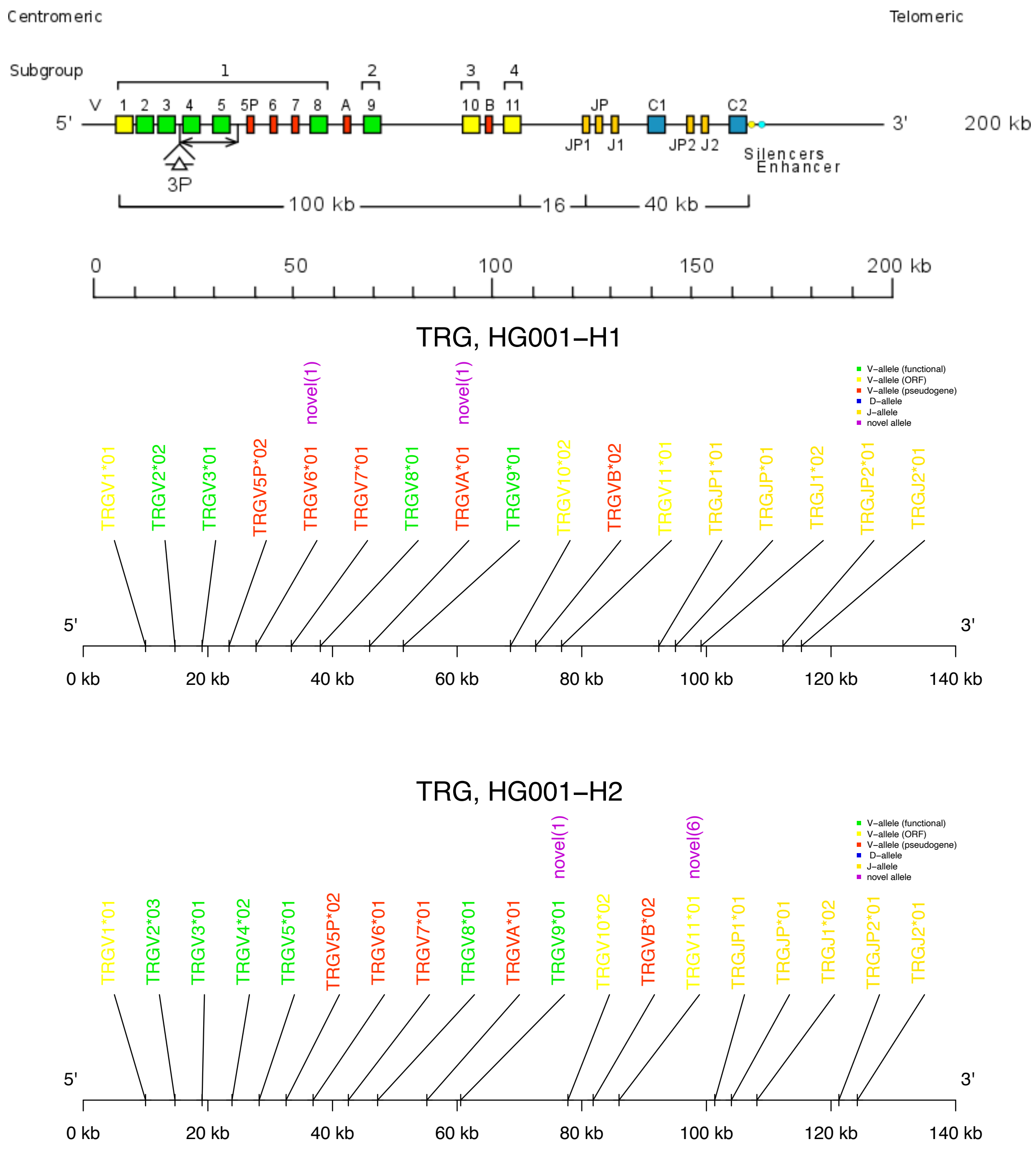


**Fig. S4.** Subfigures from top to bottom are the locus representation of TRG alleles from IMGT[32], gAIRR-annotate results of HG001-H1, and results of HG001-H2. The difference of H1 and H2 in *TRGV4* and *TRGV5* are recorded as germline variants in IMGT.

#### 7 The difference of read numbers in the deletion of HG002 chromosome 14 related to TRA/TRD alleles

**Table S5.** Number of reads aligned inside (centromeric region) and outside (telomeric region) the deletion region. Position 22,982,924 is the 3' side boundary of the deletion. The centromeric region is defined as the position 22,979,200-22,982,924. The telomeric region is defined as the position 22,982,924-22,986,400.

| Sample | centromeric region read # | telomeric region read # | drop-off rate |
|---|---|---|---|
| HG002 (son) | 944 | 1599 | 0.622 |
| HG003 (father) | 1701 | 1692 | 1.005 |
| HG004 (mother) | 2274 | 1999 | 1.138 |

#### 8 HG002's deletion on chromosome 7

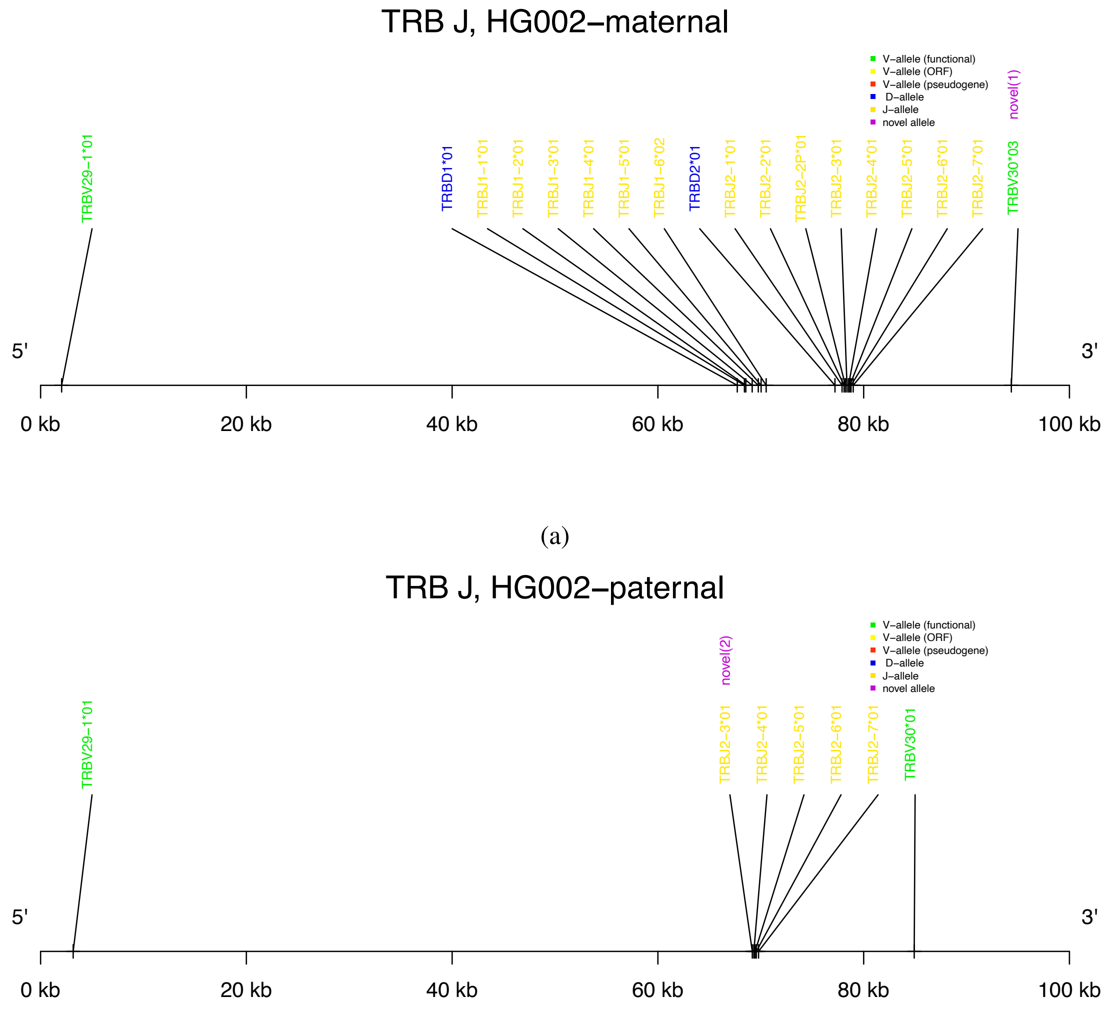


(b)

**Fig. S5.** The 10 kbp structural variation in the TRB J region of HG002. Upper: maternal haplotype, lower: paternal haplotype.

There is a 10 kb deletion on chromosome 7q34 related to TRB alleles. We align the trio capture-base short reads to GRCh38 (Supplementary Fig. S6 (a)). Although the read depth differences between HG002 and his parents are not as obvious as the deletion on chromosome 14, there are still two pieces of evidence that there is a structural variant in the area. The abrupt change indicated by the red arrow in Supplementary Fig. S6 (a) and the mate-pair alignments stretching long distances shows a deletion in one of the HG002's haplotype. It is worth mentioning that a blue arrow indicates an SNP in HG002 in Supplementary Fig. S6 (a). HG002 is C in the SNP site, while HG003 (father) is T, and HG004 (mother) is half C and half T at the same position. Lacking the C allele, the father's only haplotype in HG002 shows that the deletion is a *de novo* variation that happens to the paternal haplotype.

We choose GRCh38 rather than GRCh37 as the reference in the beta chain deletion analysis. Because the GRCh37 human genome misses the specific sequence the deletion takes place. Due to the missing segment, even HG003 and HG004 seem to have deletions in the alignment to GRCh37 (Supplementary Fig. S6 (b)). Because of the deletion in GRCh37, there are three different length insertions and one deletion reported by[31] at the site, GRCh37 chr7 position 142,494,031.

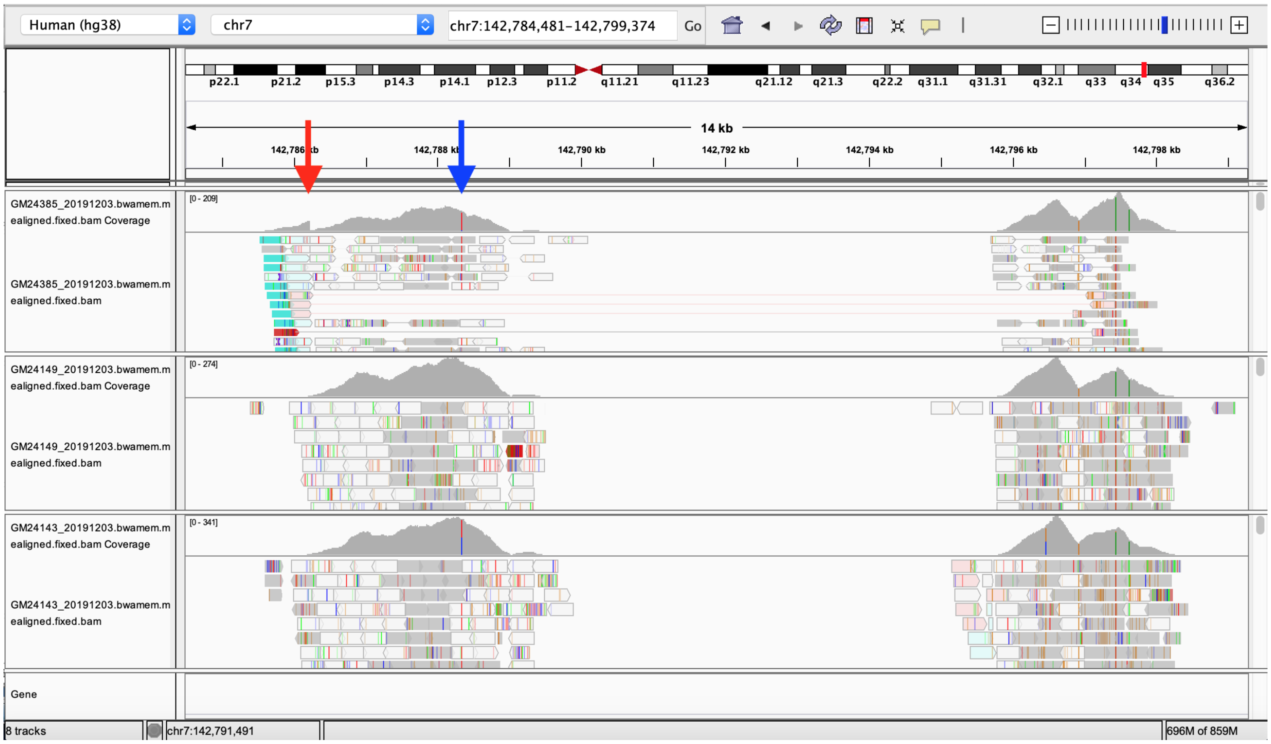


(a)

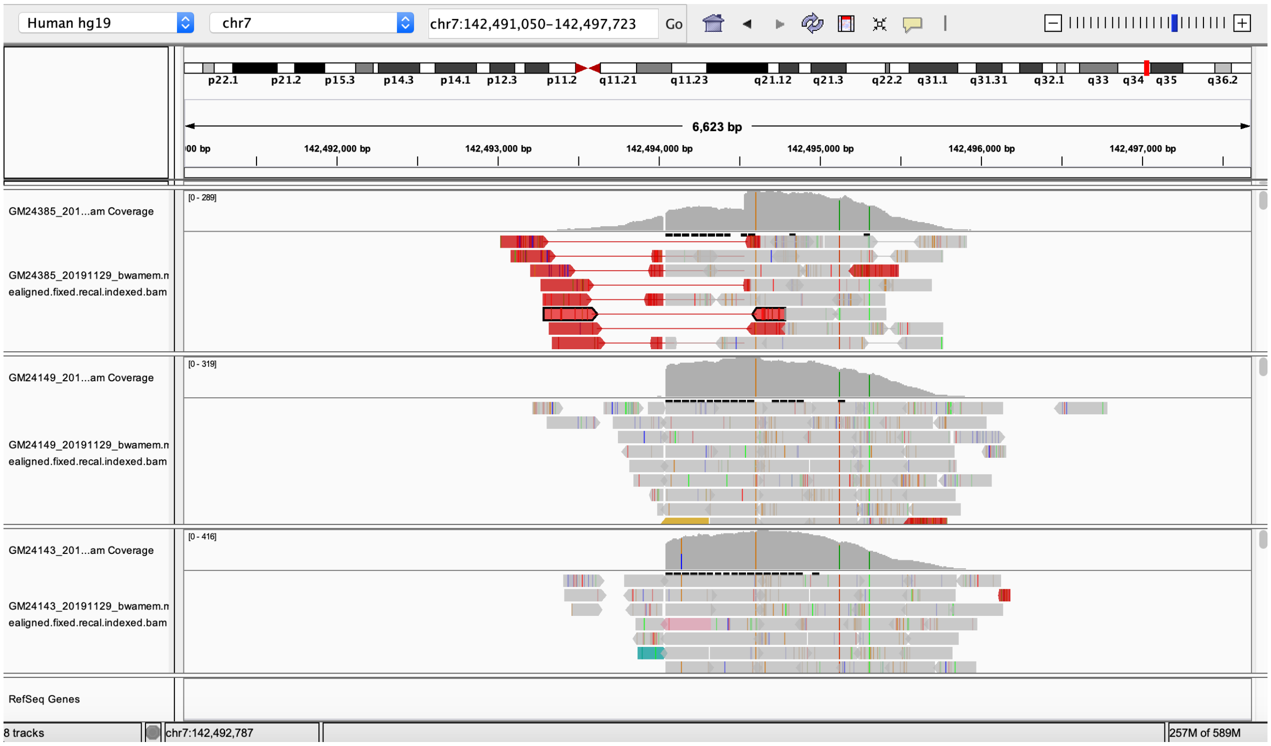


(b)

**Fig. S6.** The capture-based reads alignment of HG002 (son), HG003 (father), and HG004 (mother) to the GRCh38 chromosome 7 (a) and GRCh37 chromosome 7 (b)

#### 9 TRB locus representation of IMGT

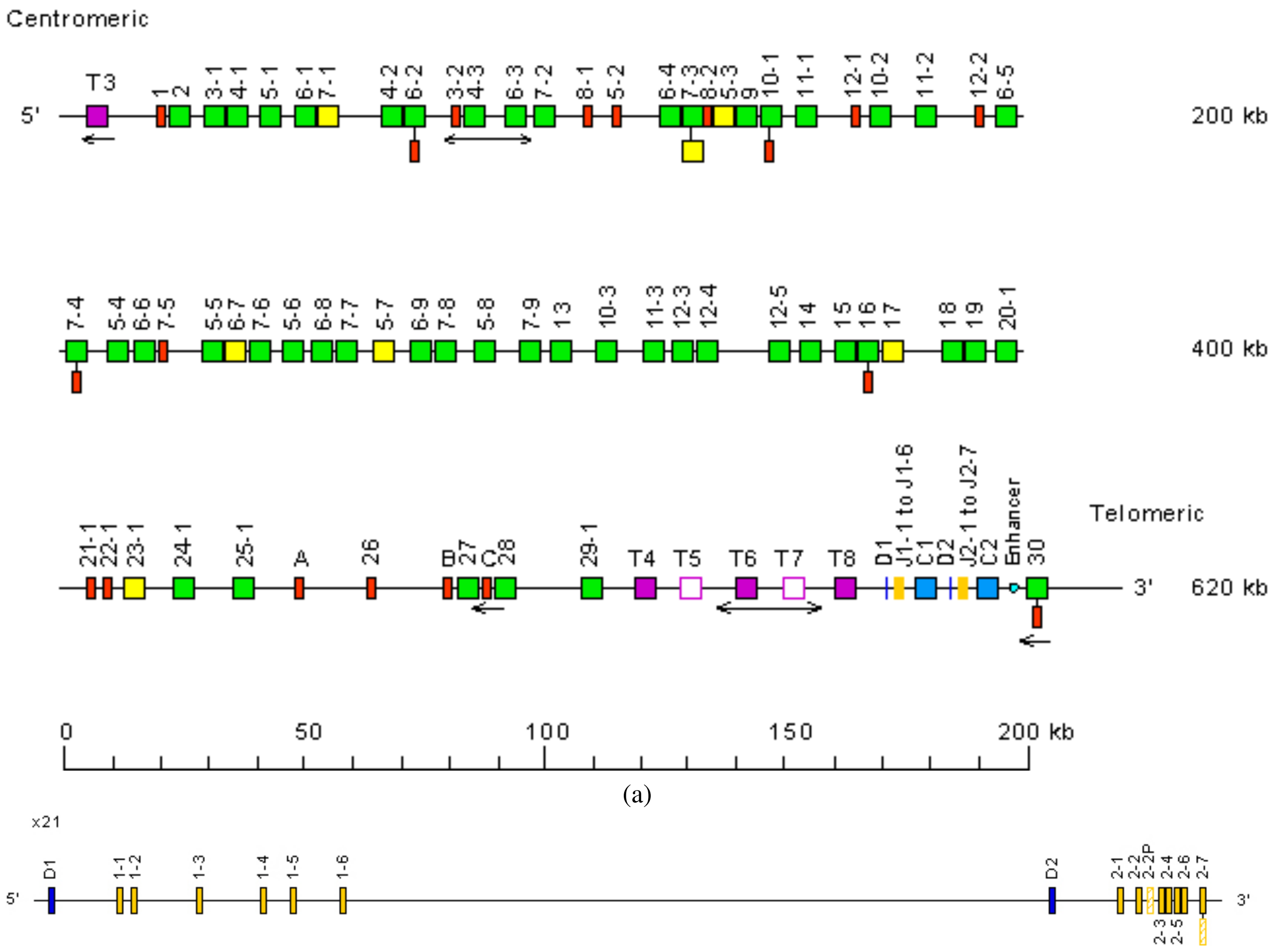


(a)

(b)

**Fig. S7.** The locus representation of TRB alleles (a) and details of TRBJ (b) downloaded from IMGT[32]

#### 10 gAIRR-annotate on IG genes

**Table S6.** Number of IG genes gAIRR-annotated from CHM13[30], GRCh37, and GRCh38. The numbers inside the brackets indicate the number of novel alleles within the total gene numbers.

| | IGH (chr14) | | | IGK (chr2) | | IGL (chr22) | | orphons | |
|---|---|---|---|---|---|---|---|---|---|
| sample | V | D | J | V | J | V | J | V | D |
| CHM13 | 132 (4) | 27 (4) | 9 | 80 (5) | 5 | 74 (19) | 7 | 96 (32) | 10 |
| GRCh37 | 116 (1) | 27 | 9 | 78 (2) | 5 | 73 (4) | 7 | 76 (23) | 10 |
| GRCh38 | 123 (3) | 27 | 9 | 78 (2) | 5 | 73 (11) | 7 | 81 (23) | 10 |
| GRCh38 chr14_KI270846v1_alt | 124 (8) | 27 | 9 | N/A | N/A | N/A | N/A | N/A | N/A |

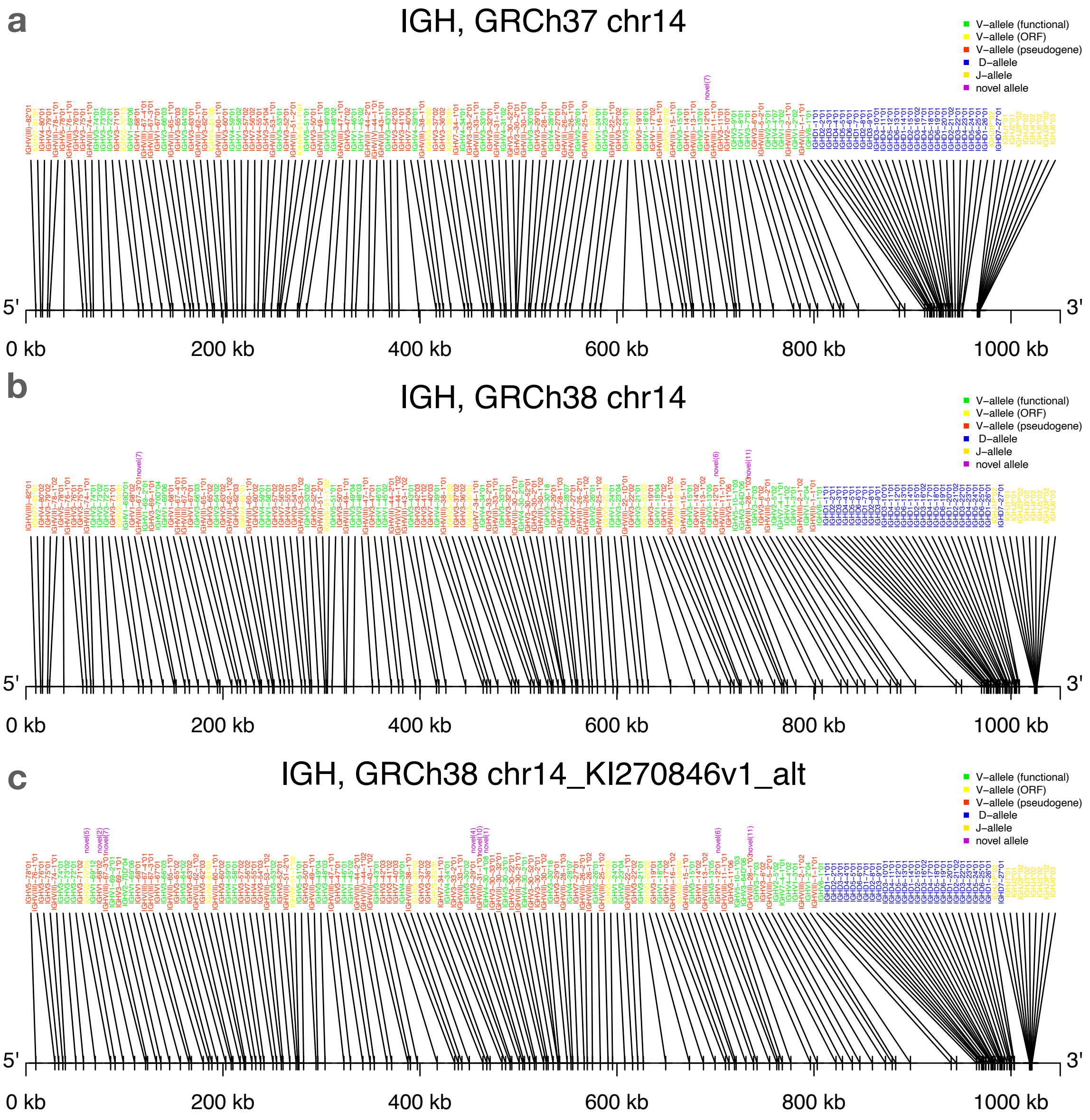


**Fig. S8.** gAIRR-annotate details on IGV alleles of the reference genomes GRCh37 and GRCh38.

#### 11 gAIRR-annotate on GRCh38's alternative contig

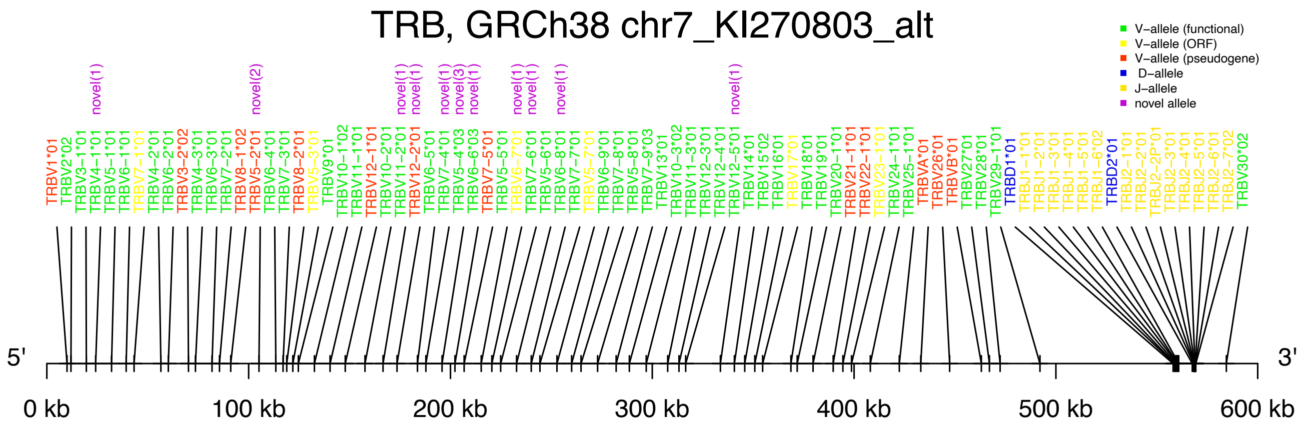


**Fig. S9.** The TR beta chain gAIRR-annotate representation of GRCh38 alternative contig chr7_KI270803v_alt.

#### 12 gAIRR-seq and gAIRR-call on primary cells

**Table S7.** The TR alleles gAIRR-call result and gAIRR-seq result of 7 GIAB RM and a Taiwanese subject's primary cell. Since the phase information is not available in gAIRR-seq and gAIRR-call, the allele numbers include both haplotypes of the samples. The total allele number, novel allele number, and read number from EBV-trasformed cell lines and primary cells are similar.

| | TRV called | | TRJ called | | gAIRR-seq result |
|---|---|---|---|---|---|
| sample | #novel | #total | #novel | #total | # of reads sequenced |
| HG001 | 26 | 186 | 3 | 86 | 280,579 |
| HG002 | 31 | 188 | 6 | 89 | 281,494 |
| HG003 | 25 | 171 | 2 | 86 | 307,312 |
| HG004 | 25 | 176 | 3 | 89 | 362,271 |
| HG005 | 29 | 166 | 3 | 86 | 327,354 |
| HG006 | 37 | 189 | 4 | 90 | 338,361 |
| HG007 | 27 | 174 | 4 | 90 | 297,186 |
| Primary cell sample | 25 | 174 | 3 | 86 | 283,691 |

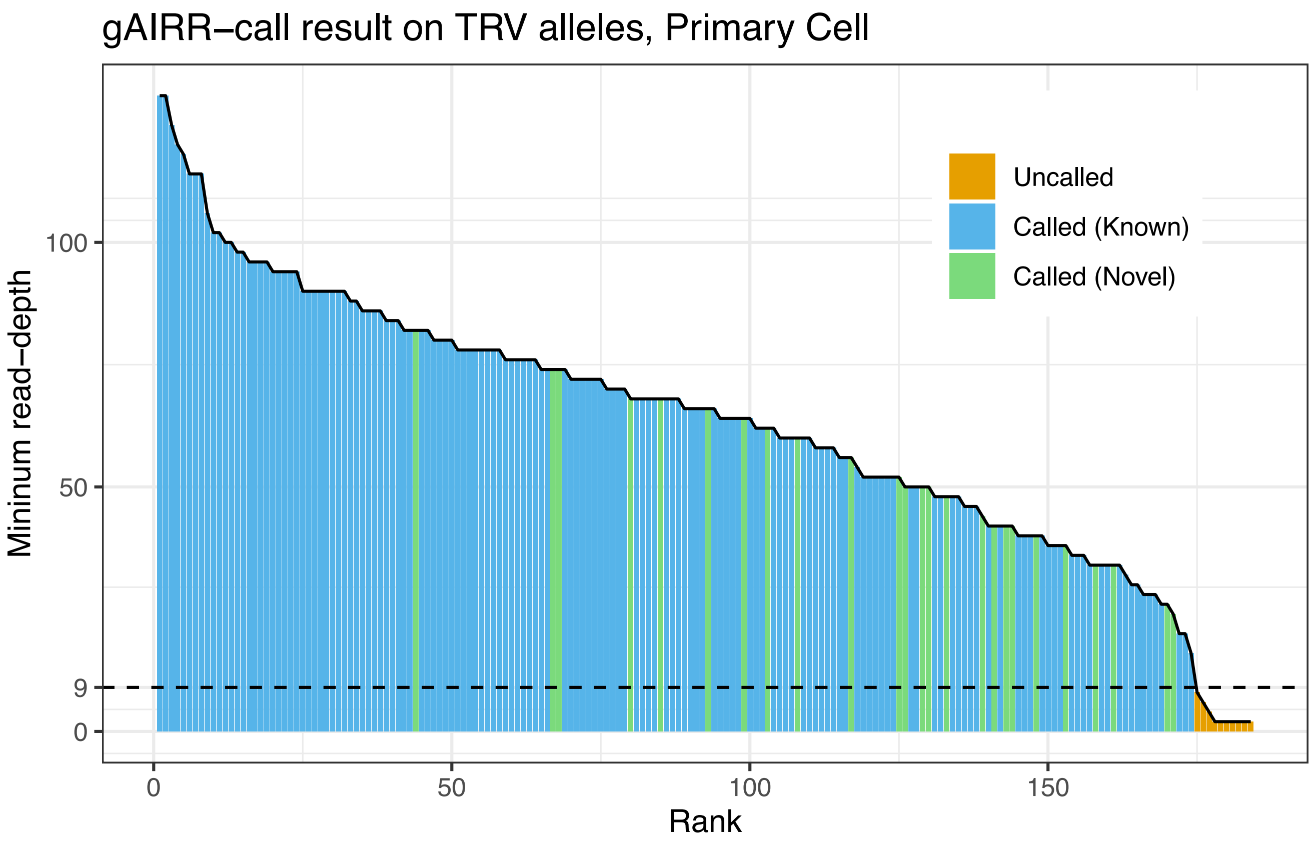


**Fig. S10.** gAIRR-call results on TRV alleles using PBMC data. The adaptive threshold is 9.

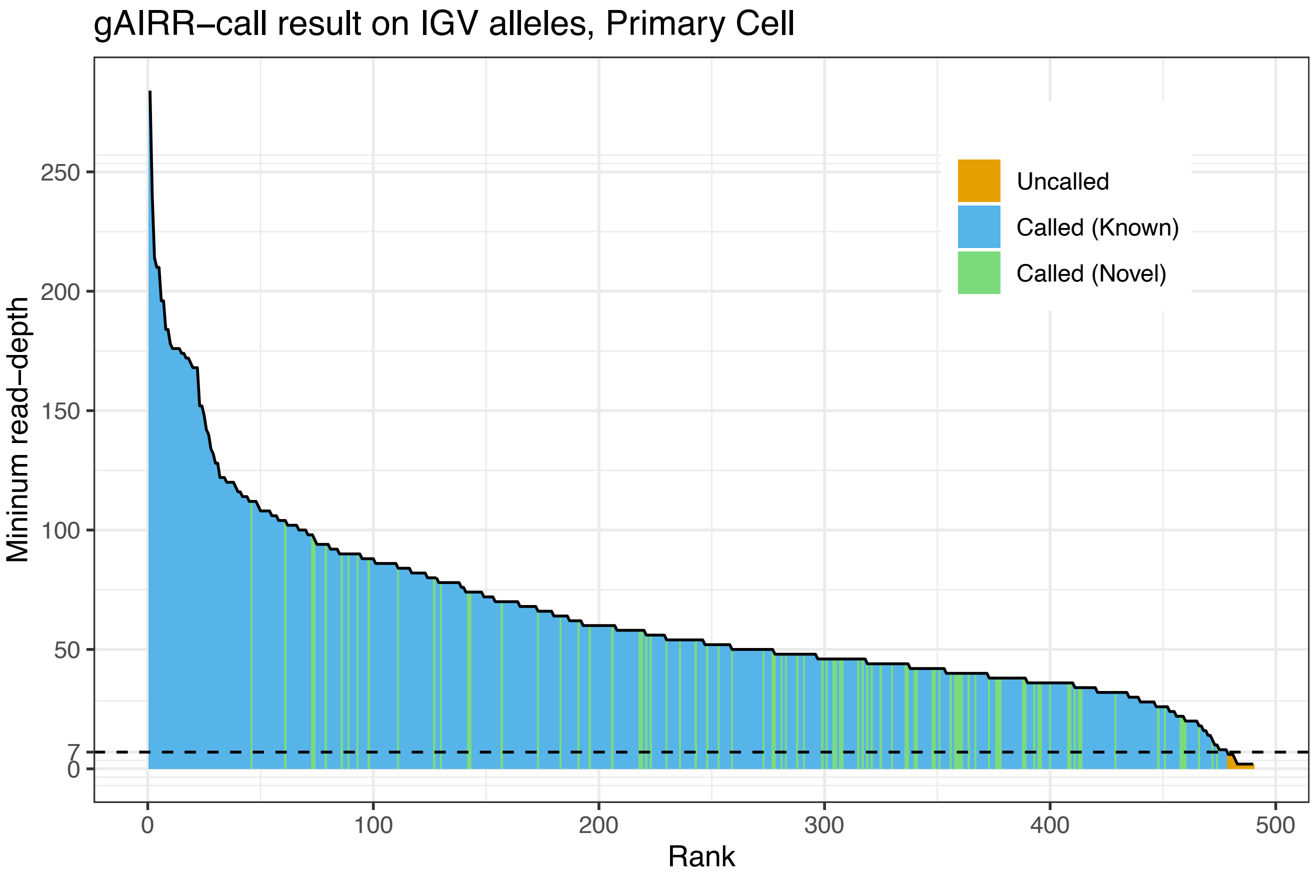


**Fig. S11.** gAIRR-call results on IGV alleles using PBMC data. The adaptive threshold is 7.

The settings of Supplementary Fig. S10 and Supplementary S11 are the same as Fig. 2b. However, there is no primary cell's personal assembly for verification. Thus, whether the called alleles are true positives is uncertain. Although the uncertainty, the result pattern of the primary cells is similar to that of HG001's EBV-transformed cell line in TRV alleles. In IGV alleles, the primary cell calling result is free from the impact of V(D)J recombination in EBV-transformed cell lines.

#### 13 AIRR allele lengths in IMGT

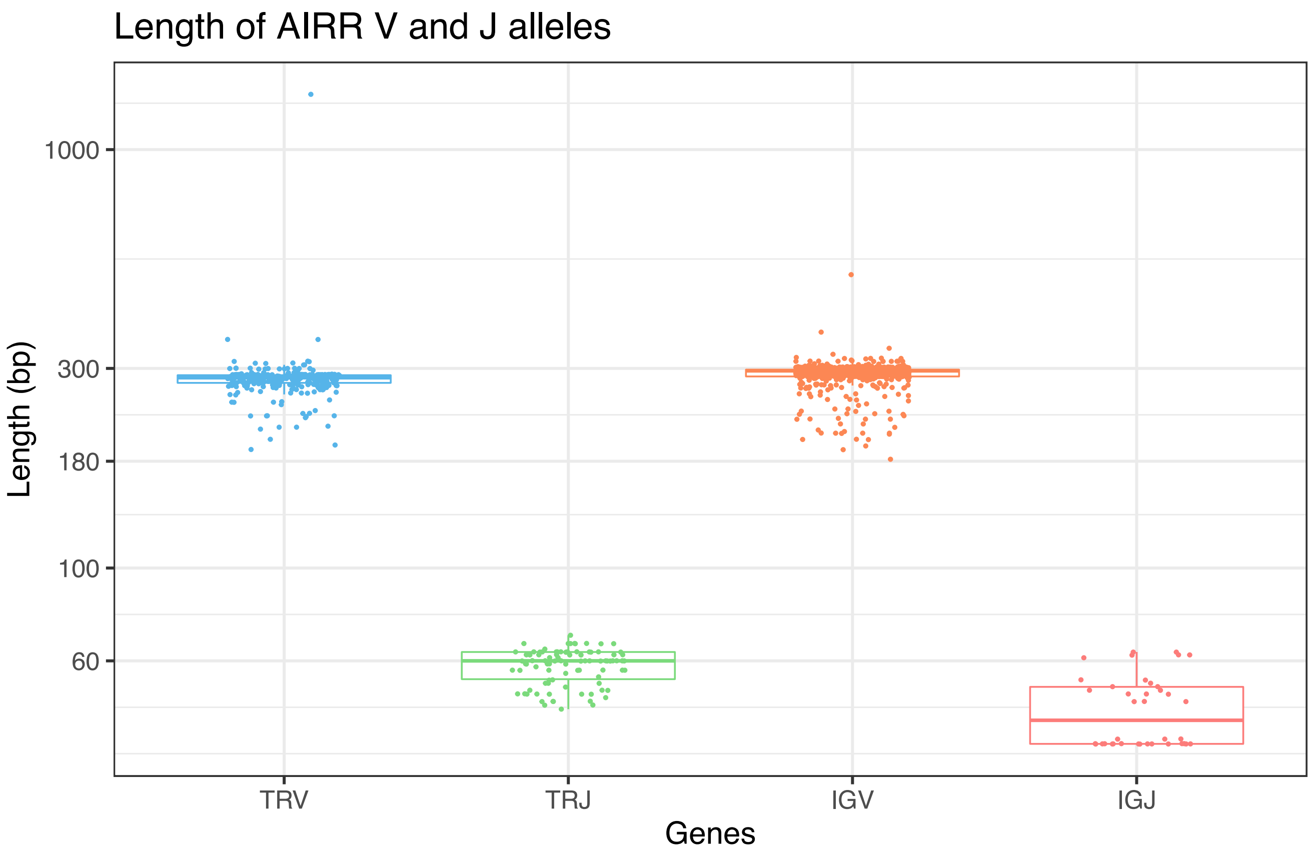


**Fig. S12.** The allele length distribution of AIRR V alleles and J alleles according to IMGT v3.1.22.

#### 14 Probe design and captured-reads representation

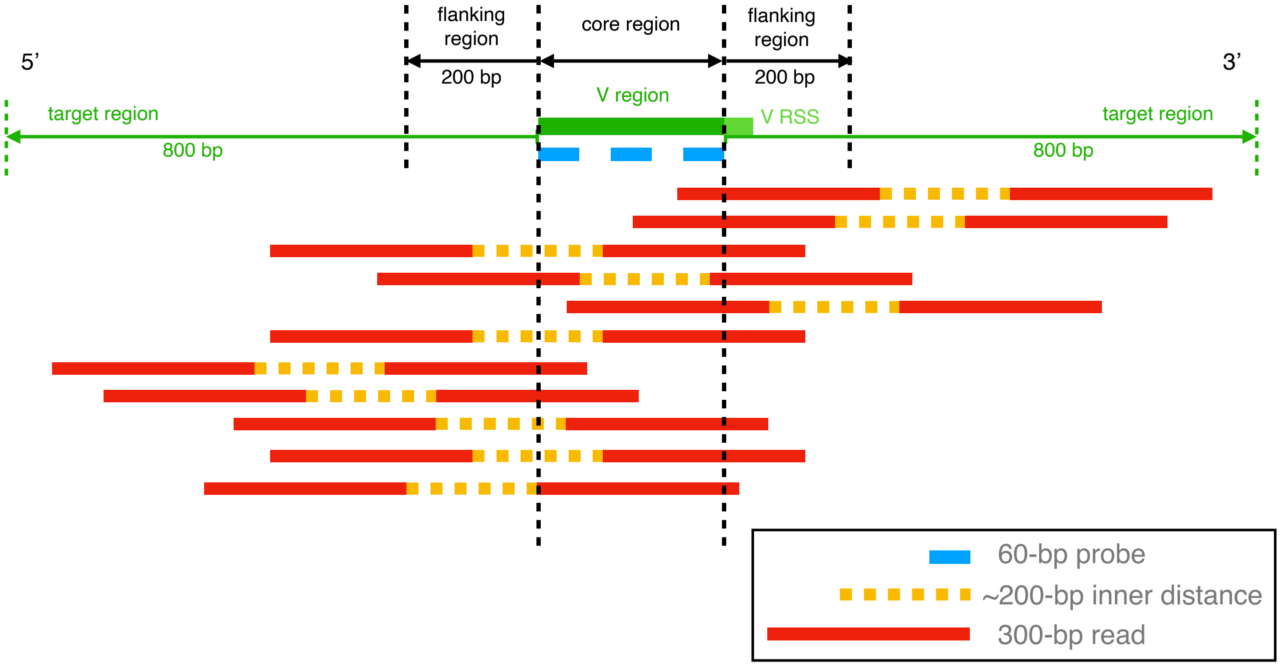


**Fig. S13.** The diagram representation of gAIRR-seq probes’ design relative to AIRR V alleles and the captured reads.

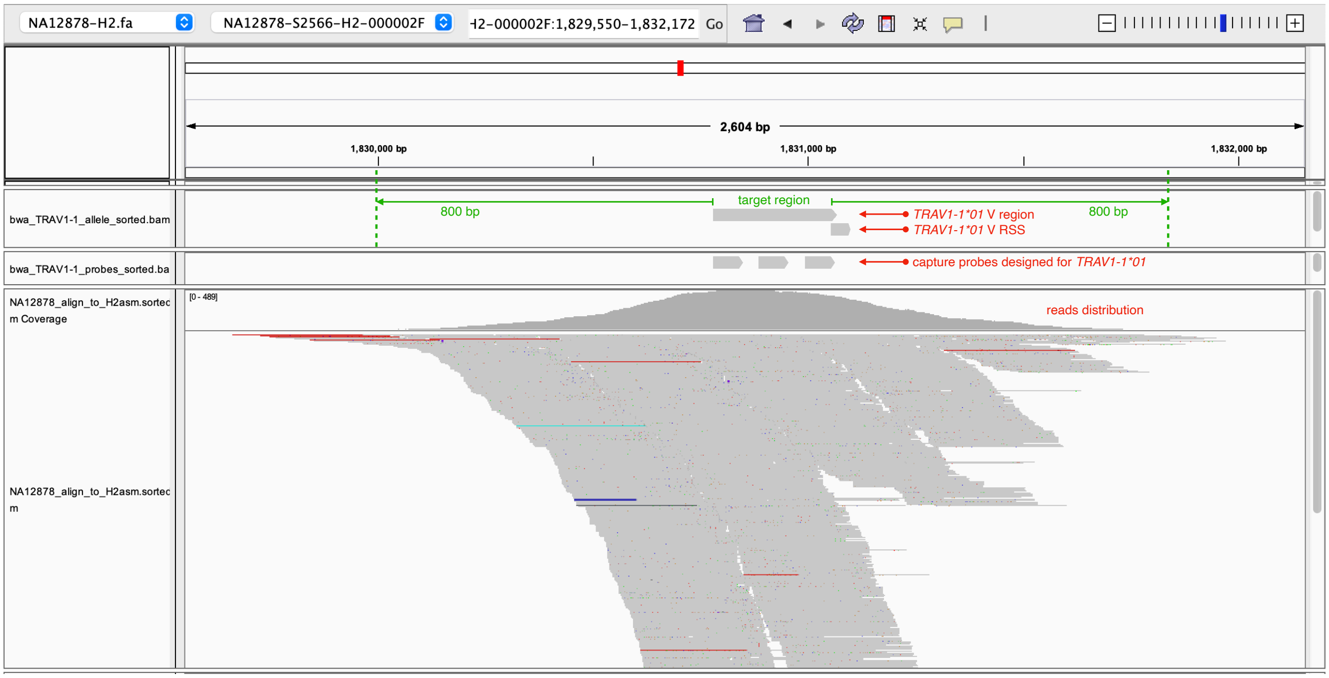


**Fig. S14.** The Integrated Genomics Viewer visualization of captured reads and probes aligned to HG001’s gene *TRAV1-1*.

In Supplementary Fig. S13, the relative positions of V alleles, V RSS, designed gAIRR-seq probes, and the reads that can be captured are shown. We also marked the range of 200 bp flanking sequences extending from the allelic region, where extended gAIRR-call and gAIRR-annotate alleles would reach. Similarly, the 800-bp range, which we defined as the target region in gAIRR-seq's on-target rate analysis is also marked. In Supplementary Fig. S14, the V allele *TRAV1-1*01*, allele's RSS, designed probes for the allele and the captured reads of HG001 are aligned to HG001's personal assembly[25]. It can be seen that most of the reads are aligned in the target region. There is a small fraction of the reads aligned outside the target region due to the variation of fragment length.

#### 15 gAIRR-annotate calls RSS in flanking sequences

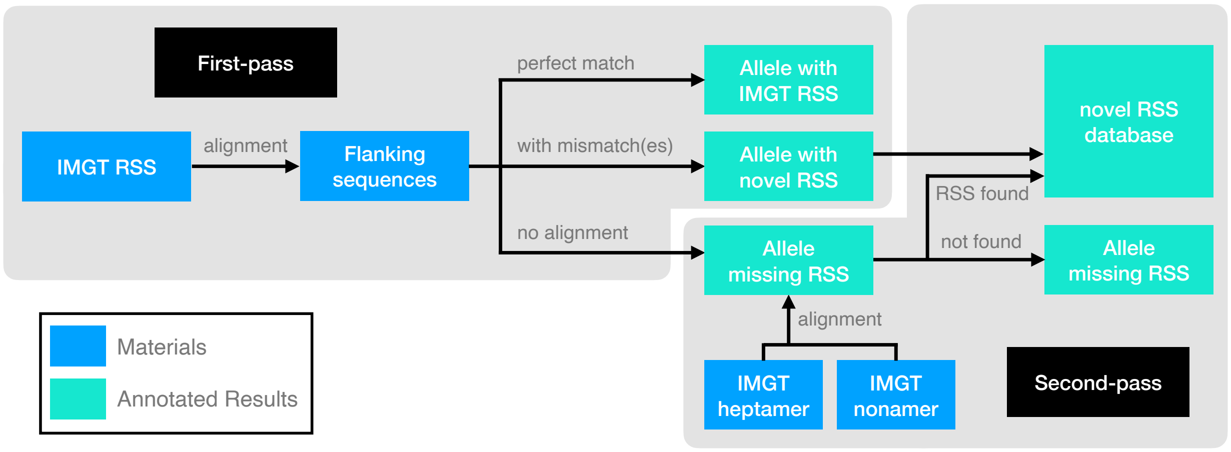


**Fig. S15.** The gAIRR-annotate pipeline to call the RSS in flanking sequences

In the first-pass, we align all IMGT RSS to the annotated flanking sequences. For an RSS aligned with mismatch(es), we define the aligned region as a novel RSS. The flanking sequences which cannot be aligned with IMGT RSS were re-aligned in the second-pass. In the second-pass, IMGT heptamers and nonamers are separately aligned to the flanking sequences, so the heptamer-nonamer pairs not recorded in IMGT can be found. Flanking sequences called by both gAIRR-call and gAIRR-annotate can be processed by this pipeline.

#### 16 Annotation files of the human reference genomes

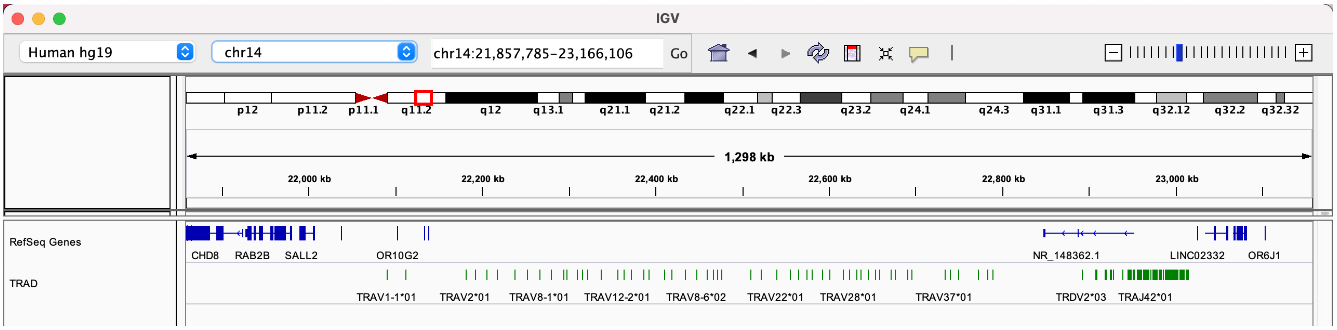


**Fig. S16.** Loading the TR alpha/delta chain annotation file hg19_TRAD.bed into the Integrated Genomics Viewer[39].

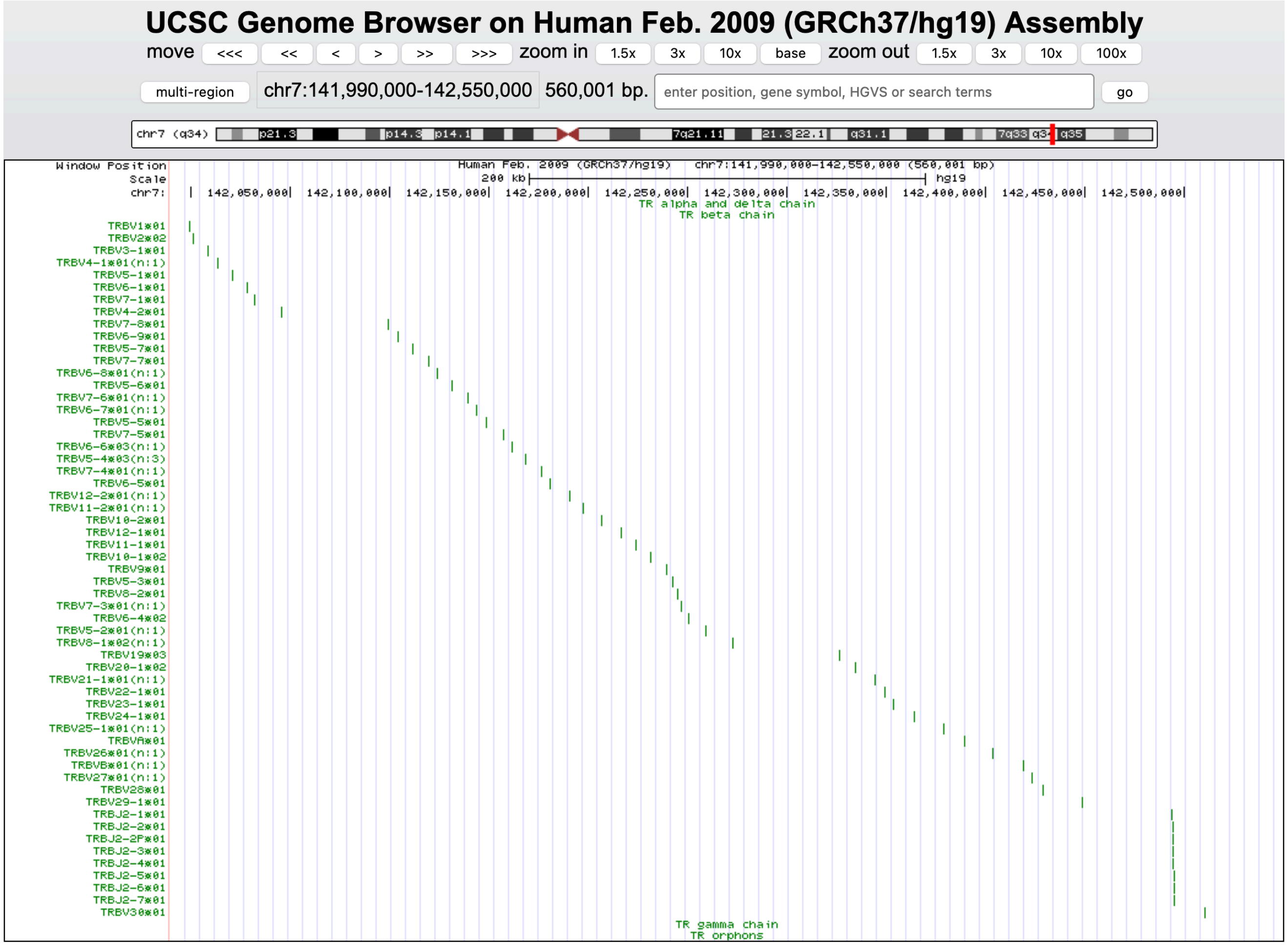


**Fig. S17.** Loading the TR beta chain annotation file hg19_TRB.bed into the UCSC Genome Browser[40].

Under the directory *supplementary_files/reference_genome_annotation/* in the GitHub link `https://github.com/maojanlin/gAIRRsuite` are the gAIRR-annotated bed files of the human reference genomes. In the bed files, the novel alleles will be indicated by brackets after allele names. The number after 'n:' in the brackets is the number of mismatches of the novel allele. There are four bed files recording the allele positions of GRCh37's four TR loci. Similarly, five bed files are for GRCh38's four TR loci and one locus on the alternative contig. Users can easily load the bed files into visualization tools such as Integrated Genomics Viewer[39] (Supplementary Fig. S16) or UCSC Genome Browser[40] (Supplementary Fig. S17).
